# Aerobic denitrification as N_2_O source in microbial communities

**DOI:** 10.1101/2023.06.14.544945

**Authors:** Nina Roothans, Minke Gabriëls, Martin Pabst, Mark C. M. van Loosdrecht, Michele Laureni

**Affiliations:** Delft University of Technology, Mekelweg 5, 2628 CD Delft, the Netherlands

## Abstract

Nitrous oxide (N_2_O) is a potent greenhouse gas of primarily microbial origin. Aerobic and anoxic emissions are commonly ascribed to nitrification and denitrification, respectively. Beyond this established dichotomy, we quantitatively prove that heterotrophic denitrification can significantly contribute to aerobic nitrogen turnover and N_2_O emissions in complex microbiomes exposed to frequent oxic/anoxic transitions. Planktonic, nitrification-inhibited denitrifying enrichments respired over a third of the influent organic substrate with nitrate at high oxygen concentrations. N_2_O accounted for up to one quarter of the aerobically respired nitrate. The constitutive detection of all denitrification enzymes in both anoxic and oxic periods highlight the selective advantage offered by metabolic preparedness in dynamic environments. We posit that aerobic denitrification and associated N_2_O formation is currently underestimated in dynamic microbial ecosystems.

## Introduction

Nitrous oxide (N_2_O) is today’s third most important greenhouse gas and the main stratospheric ozone-depleting substance^1^. Globally, the majority of N_2_O originates from biological conversions in natural, managed, and engineered ecosystems^2^, such as oceans^3^, agricultural soils^4^, and wastewater treatment plants^5^. N_2_O emissions from anthropogenic activities are projected to reach 11.5 Tg N yr^-1^ in 2050, doubling their amount in 2000, if no mitigation action is taken^1,2^. Robust emission control strategies strongly rely on our knowledge of the microbiology underlying N_2_O turnover.

N_2_O is a metabolic by-product of nitrification, the aerobic oxidation of ammonium (NH_4_^+^) to nitrite (NO_2_^-^) and nitrate (NO_3_^-^), and an obligate intermediate of denitrification, the multi-step reduction of NO_3_^-^ to dinitrogen gas (N_2_). Conventionally, nitrification and denitrification are considered to dominate N_2_O emissions in the presence and absence of O_2_, respectively^3,4,6^. Oxygen is known to regulate the expression and inhibit the activity of denitrifying enzymes^7^. As most known denitrifiers are facultative aerobes, the more energetically and kinetically favourable aerobic respiration is expected to be prioritized over denitrification in the presence of O_2_^8^. Thus, the aerobic contribution of denitrification is generally neglected in soils^9–11^, oceans^3,12^, and wastewater treatment systems^13–15^. The impact of O_2_ differs per denitrifying enzyme, with the nitrous oxide reductase (NosZ) - the only known biological N_2_O sink - appearing to be the most sensitive enzyme^7,16,17^. Starting from the seminal work of Robertson & Kuenen^18^, the occurrence of denitrification under high oxygen concentrations has been documented in pure culture studies (as reviewed by Chen & Strous^8^). The ecological significance of denitrification in aerobic N_2_O formation remains to date largely unknown.

*Sensu stricto*, aerobic denitrification is the simultaneous respiration of O_2_ and nitrogen oxides^19–21^. Aerobic denitrifying bacteria, including *Paracoccus denitrificans, Alcaligenes faecalis* and multiple *Pseudomonas* species, have been successfully isolated mainly from ecosystems exposed to fluctuating O_2_ levels such as soils, sediments and activated sludge^18,21– 23^. Aerobic denitrification rates are generally lower than the anoxic rates, yet are suggested to provide an ecological advantage in dynamic environments^8,18,21,24^. At the same time, these results are based on a limited number of isolates characterized primarily under continuous aeration, inherently hindering their extrapolation to complex microbiomes in dynamic O_2_ environments. Central challenges in open ecosystems are the co-occurrence of nitrification as potential confounding aerobic N_2_O source, and the development of anoxic micro-niches in microbial aggregates^25,26^. Only three studies quantified aerobic denitrification in natural communities, namely in soil bacteria extracted by density-gradient centrifugation^26^ and intact sea sediments^20,25^. All authors experimentally showed nitrification to be negligible, yet anoxic niches could not be completely excluded. Gao et al.^25^ even observed a marked decrease in aerobic NO_x-_ respiration upon vigorous stirring, possibly resulting from the disruption of anoxic micro-niches. The unresolved question remains to which extent the long-overlooked aerobic denitrification contributes to overall nitrogen turnover in dynamic ecosystems.

We enriched for two heterotrophic denitrifying communities co-respiring O_2_ and NO_3_^-^ under alternating oxic/anoxic conditions to quantitatively resolve the role of aerobic denitrification in mixed communities. Fully aerated planktonic cultures were employed to exclude anoxic micro-niches, while continuous allylthiourea (ATU) addition ensured full suppression of nitrification. The metabolic potential of and actual labour-division within the enrichments were characterized by metagenomic and metaproteomic analysis. To the best of our knowledge, this is the first study quantitatively proving that aerobic denitrification can significantly contribute to nitrogen-turnover and N_2_O emissions in microbial communities with time-varying oxygen availabilities; and suggests that nitrifiers’ contribution to aerobic N_2_O emissions may currently be overestimated.

## Results

### 1. Stable denitrifying cultures under alternating oxygen availability

Two planktonic denitrifying microbial communities were enriched under alternating anoxic and fully aerobic conditions. A mixture of volatile fatty acids (acetate, propionate, butyrate) served as carbon and energy source, and NO_3_^-^ as electron acceptor. All dissolved substrates were continuously provided (Supplementary Table S1). The O_2_ supply was controlled to ensure a 1:2 ratio of anoxic to oxic time, split in 4 (R_4_) and 32 (R_32_) cycles per day. Fully anoxic conditions were ensured by continuous N_2_ sparging. In the oxic phase, dissolved oxygen was maintained above 6 mg/L (>75% air saturation), and both NO_3_^-^ and O_2_ served as electron acceptors. Continuous supply of allylthiourea (ATU) ensured complete suppression of nitrification, as confirmed by the absence of ammonium oxidation activity (day 61, Supplementary Figure S3) and nitrification genes in the recovered metagenomes (Figure 1).

**Figure 1.**
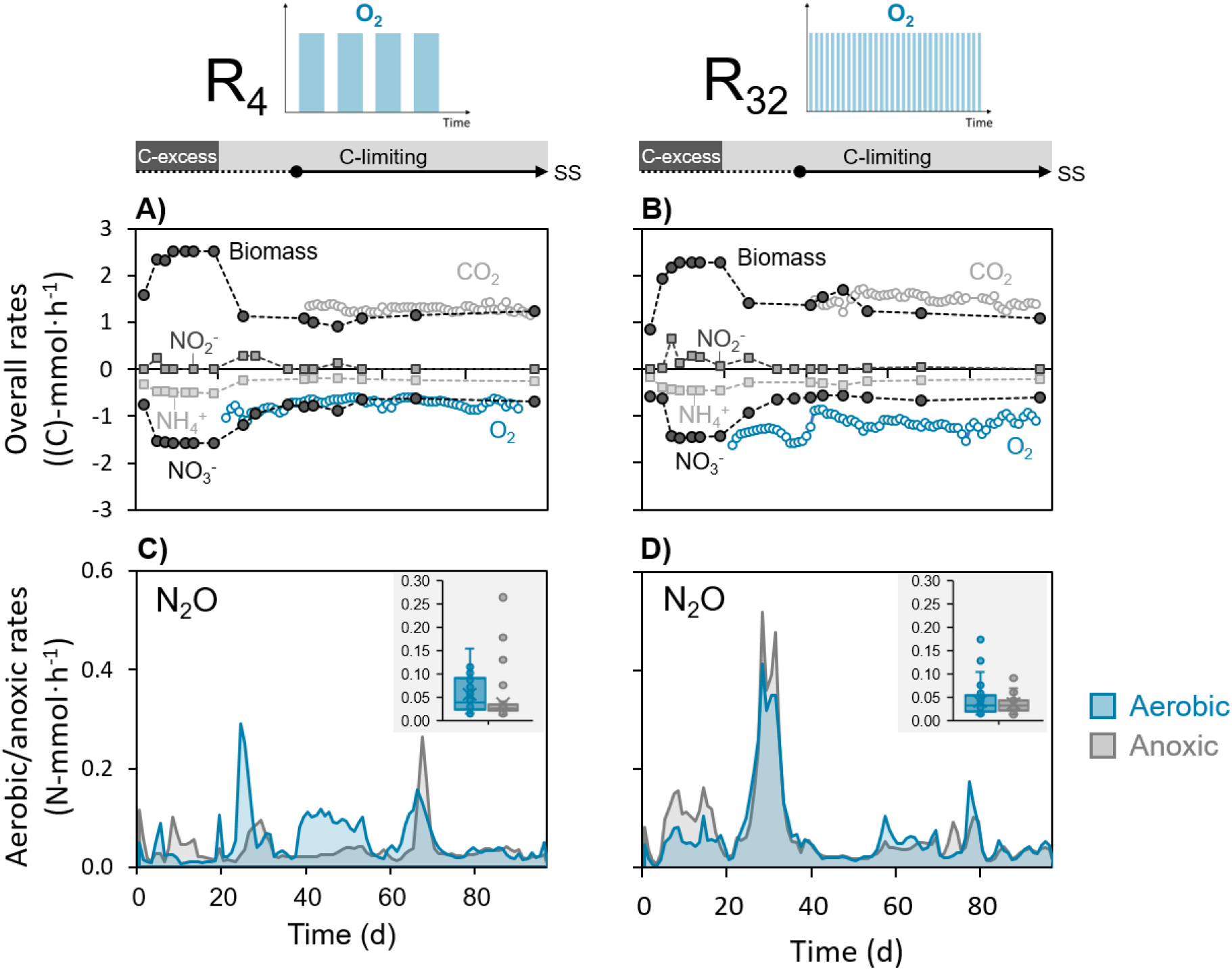
Conversion rates (mmol·h^-1^) in the low-(R4) and high-frequency (R32) oxic/anoxic cycling reactors over the entire operational period. Prior to the target carbon limiting conditions, the reactors were started up for 27 days under carbon excess. The steady-state (SS), was reached on day 37 and maintained for over two months (equivalent to 30 generation times). Negative rates represent consumption while positive rates represent production. **Panels A and B:** overall (*i*.*e*. combined oxic and anoxic) NO_3_^-^, NH_4_^+^ and O_2_ consumption, and NO_2_^-^, CO_2_ and biomass production rates (in C-mmol·h^-1^). The latter was calculated from the NH_4_^+^ consumption rate. For consistency, an “overall” O_2_ consumption rate was calculated by averaging its aerobic consumption over the entire cycle duration. Error bars of all rates are smaller than the points and represent the standard deviation of triplicate samples (nitrogen substrates) or of daily averages of continuous measurements (CO_2_ and O_2_). **Panels C and D:** Daily average N_2_O production rates (N-mmol/h) during the aerobic and anoxic phases. Insert: boxplots summarize the daily N_2_O emission rates (N-mmol/h) in both phases during the steady-state period.

After a start-up period of 21 days, the reactors were run for 76 days (equivalent to 38 volume changes) under carbon-limiting conditions with a dilution rate of 0.02 h^-1^. The operational steady-state was reached after day 37, as confirmed by constant overall substrates and products conversion rates (Figure 1, panels A-B). These overall rates represent the weighted average of the oxic and anoxic rates within one cycle (Supplementary eq. S9). For consistency, an “overall” consumption rate was also calculated for O_2_, by averaging its aerobic consumption over the entire cycle duration (Supplementary Section 2). The overall NH_4_^+^, CO_2_, organic carbon, and biomass conversion rates (Supplementary Table S1), as well as the resulting stoichiometric yields (Table 1), were comparable between the two reactors. The enrichments differed only in terms of the overall NO_3_^-^ and O_2_ yields (Table 1). Over the combined oxic and anoxic periods, 56±4% and 39±4% of the total catabolic electron flow was used for NO_3_^-^ reduction in R_4_ and R_32_ respectively, with the remaining being used for O_2_ reduction (Supplementary Table S2). NO_2_^-^ and NO accumulation was absent or negligible during the entire steady-state period. The carbon and electrons balances closed, further confirming that all involved substrates and products were measured, and supporting N_2_O and N_2_ as the primary products of NO_3_^-^ reduction (Table 1).

**Table 1.** Average overall (*i*.*e*. combined oxic and anoxic) steady-state stoichiometric yields and carbon and electron balances in the low-(R4) and high-frequency (R32) reactors. The standard deviations were calculated from the standard deviation of the consumption and production rates (Supplementary Table S1) using linear error propagation (Supplementary eq. S3).

| Reactor | Y<sub>N2O/NO3-</sub><br>(Nmol/Nmol) | Y<sub>X/S</sub><br>(Cmol/Cmol) | Y<sub>NO3-/S</sub><br>(Nmol/Cmol) | Y<sub>O2/S</sub><sup>a</sup><br>(mol/Cmol) | Y<sub>CO2/S</sub><br>(Cmol/Cmol) | C bal<br>(%) | e<sup>-</sup> bal<br>(%) |
| --- | --- | --- | --- | --- | --- | --- | --- |
| R<sub>4</sub> | 0.07 ± 0.04 | 0.44 ± 0.04 | 0.30 ± 0.04 | 0.29 ± 0.03 | 0.53 ± 0.03 | 103 ± 5 | 101 ± 8 |
| R<sub>32</sub> | 0.07 ± 0.04 | 0.46 ± 0.06 | 0.20 ± 0.01 | 0.40 ± 0.05 | 0.51 ± 0.05 | 103 ± 8 | 100 ± 8 |

<sup>a</sup> For consistency, the O<sub>2</sub> respiration yield is the “overall” (*i.e.* combined oxic and anoxic) instead of the aerobic yield.

### 2. Comparable average aerobic and anoxic N_2_O production rates

The aerobic and anoxic N_2_O production rates remained highly variable throughout the entire operation (Figure 1, panels C-D), despite both systems being at apparent steady-state (after day 37). The daily average N_2_O emission rates fluctuated between 0.02 and 0.16 N-mmol·h^-1^ in the two systems. The average N_2_O production rate in R_4_ was higher in the oxic than in the anoxic phase (0.057±0.037 *vs*. 0.037±0.039 N-mmol/h), while these were nearly identical in R_32_ (0.042±0.029 *vs*. 0.038±0.019 N-mmol/h).

The high aerobic N_2_O production implies that denitrification was active at fully aerobic conditions (> 6 mg O_2_/L). The aerobic and anoxic NO_3_^-^ consumption rates were estimated based on the aerobic and anoxic organic substrate and oxygen consumption, CO_2_ production and N_2_O accumulation rates, and the electron balances in each phase (Supplementary Section 2). The estimated aerobic NO_3_^-^ consumption rates were only 2.4- and 7.7-fold lower than the anoxic rates in R_4_ and R_32_, respectively. This is equivalent to 36±7% and 11±11% of the total aerobic electron flow in each reactor. The fraction of NO_3_^-^ emitted as N_2_O during aeration was estimated to be 12±8% (R_4_) and 24±29% (R_32_).

### 3. Denitrifiers-enriched microbial communities

Long-read sequencing of the whole community DNA (day 68) yielded over 2 and 0.5 million reads with N50 of 5.9 and 6.2 kb for R_4_ and R_32_, respectively, after quality filtering and trimming. Reads assembly resulted in 2747 and 2002 contigs with N50 of 151 and 240 kb. After binning, we recovered a total of 21 (R_4_) and 18 (R_32_) high-quality metagenome-assembled genomes (MAGs) with over 90% completeness and under 5% contamination (Supplementary Table S4 and S5). The top 10 most abundant high-quality MAGs accounted for 78% (R_4_) and 57% (R_32_) of the mapped reads normalized to the corresponding MAG length (Figure 2). Low-abundant high-quality and all medium-quality MAGs (< 90% completeness and > 5% contamination) were grouped into “others”. Low-quality bins (<70% completeness or >10% contamination) were grouped with the unbinned fraction, accounting for 18% (R_4_) and 26% (R_32_) of the community. MAG-based taxonomic analysis revealed two distinct communities, both dominated by the Proteobacteria phylum (Supplementary Table S4 and S5). R_4_ was co-dominated by members of the *Denitromonas* (Gammaproteobacteria) and *Wagnerdoeblera* (Alphaproteobacteria) genera (Figure 2). In R_32_, the two most abundant MAGs belonged to the *Castellaniella* genus (Gammaproteobacteria).

**Figure 2.**
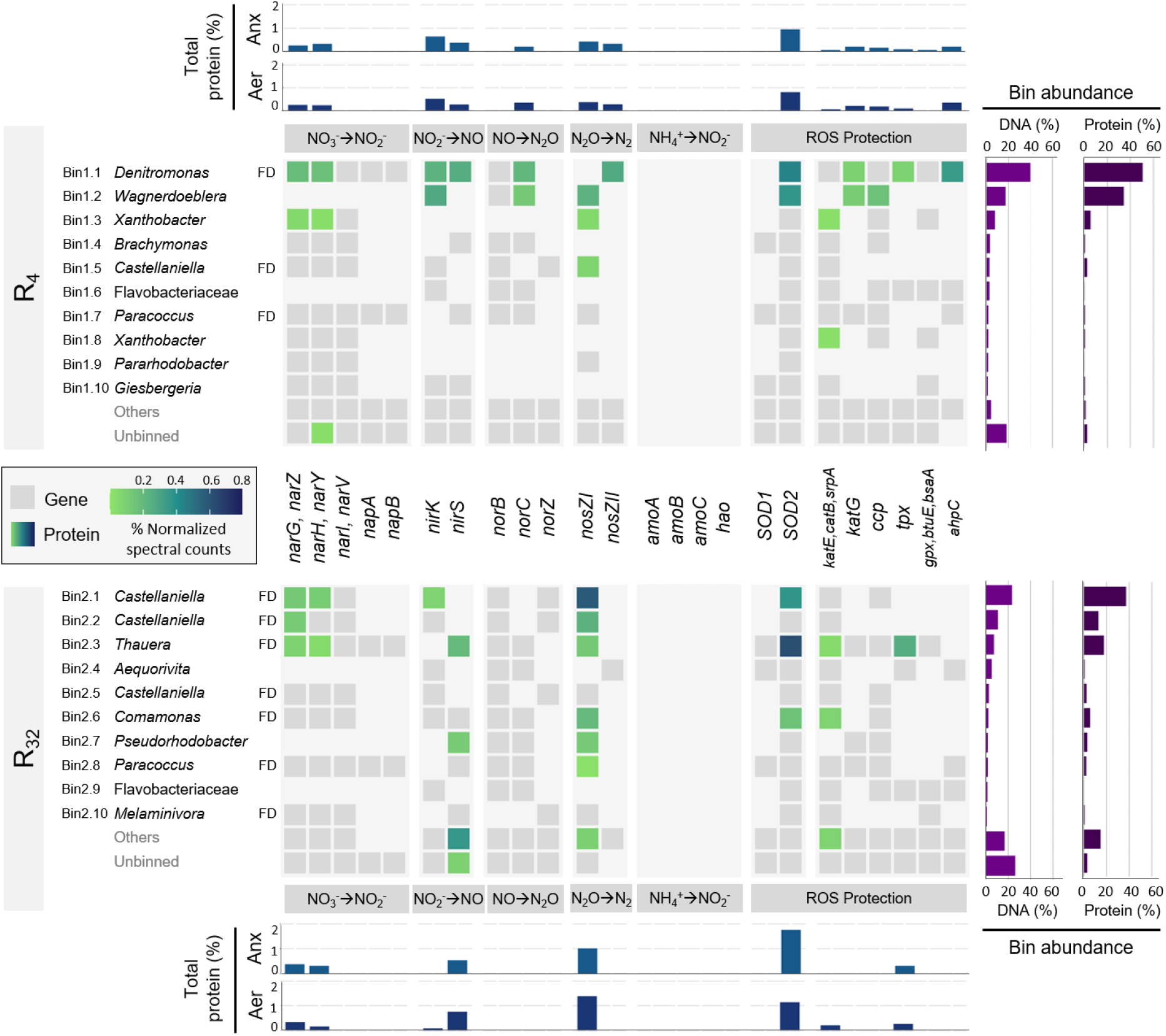
Genomic and proteomic profiles of the top 10 most abundant high-quality MAGs in both enrichments. Gene presence and protein expression in high-quality MAGs (completeness > 90% and contamination < 5%) in the low-(R4) and high-frequency (R32) reactors (top panel – R4 – and lower panel – R32), with their respective taxonomic classification at genus (or family if unclassified genus) level. Full denitrifying organisms, with genes encoding for all denitrification steps, are highlighted (FD). Low-abundant high-quality and all medium-quality MAGs (< 90% completeness and > 5% contamination) were grouped into “others” and low-quality bins (< 70% completeness and > 10% contamination) were grouped with the unbinned fraction. The presence of genes (grey tiles) and the abundance of their corresponding protein under aerobic conditions (coloured tiles) are represented for denitrification (NO_3_^-^ → N2), nitrification (NH_4_^+^ → NO_2_^-^) and protection against reactive oxygen species (ROS). The abundance of each protein was determined from peptide spectral sequence counts. **Right bar charts:** total relative abundance of each MAG in the metagenome (based on relative reads alignment normalized to the corresponding MAG length) and the metaproteome (summed relative abundance of normalized spectral counts of peptides matching to predicted proteins in each MAG). **Top/bottom bar charts:** total relative abundance of each protein in the aerobic and anoxic phases (summed relative abundance of normalized spectral counts).

All high-quality MAGs contained at least one gene of the denitrification pathway, and full denitrifiers dominated the community in R_32_ (Figure 2 and Supplementary Section 5). The membrane-bound NO_3_^-^ reductase gene (*narGHI*) was annotated in most MAGs, while only a few also possessed the periplasmic reductase gene (*napAB*). Most MAGs had either a Cu-type (*nirK*) or *cd1*-type (*nirS*) NO_2_^-^ reductase gene, with some possessing both. Overall, the cytochrome c-dependent nitric oxide reductase genes (*norBC*) were more frequent than the quinol-dependent reductase genes (*norZ*). *norZ* in members of the *Castellaniella* genus were always accompanied with an additional *norB* gene. The N_2_O reductase gene (*nosZ*) was widespread in both reactors, and was dominated by the clade I type. No subunits of the ammonia monooxygenase (*amoABC*) and hydroxylamine oxidoreductase (*hao*) genes were found. Also, the *nrfAH* genes, catalysing the dissimilatory reduction of NO_2_^-^ to NH_4_^+^, were essentially absent in the MAGs (Supplementary Section 5). All denitrifying MAGs also contained the genes encoding the O_2_-reducing terminal oxidases (complex IV) (Supplementary Section 5), and enzymes protecting against reactive oxygen species (ROS), including superoxide dismutases (SOD) and catalases/peroxidases (Figure 2).

### 4. Highly comparable anoxic and aerobic proteomic profiles

Shotgun metaproteomics of the steady-state enrichments (day 68) revealed the aerobic and anoxic expression of key denitrification and ROS-protecting enzymes by each MAG (Figure 2 and Supplementary Section 5). Over 70% (R_4_) and 50% (R_32_) of the detected total peptide intensity (peak area) uniquely matched with proteins predicted from the respective metagenomes. A total of 750/849 and 724/576 proteins of R_4_ and R_32_ (aerobic/anoxic) were identified by at least two unique peptides. The protein-based relative abundance of most MAGs was consistent with their genome-based abundance (Figure 2, right bar charts). The contribution to the overall proteome of the unbinned and others fraction combined, accounting for 22% and 43% of the metagenomes, was only 4% and 23% for R_4_ and R_32_, respectively.

The overall and MAG-specific relative abundances of the detected denitrification enzymes was highly comparable between the oxic and anoxic phase in each enrichment (Figure 2 and Supplementary Section 5). The catalytic subunits of the membrane-bound NO_3_^-^ reductase (NarG), Cu-type (NirK) or *cd1*-type (NirS) NO_2_^-^ reductase and N_2_O reductase (NosZ) were consistently present. NosZ I and NosZ II were both expressed in R_4_, while only NosZ I was detected in R_32_. In R_4_, the two most abundant MAGs (bin1.1 and bin1.2) accounted for most of the expressed denitrification proteins. On the contrary, in R_32_, lower abundant MAGs significantly contributed to the expression of NirS and NosZ. Moreover, NirS was the dominant type of NO_2_^-^ reductase detected in R_32_. The membrane-bound cytochrome c-(cNor) and quinol-dependent (qNor) NO reductases were not detected, nor was the periplasmic NO_3_^-^ reductase (NapAB) (Figure 2). With respect to oxygen, the abundance of the superoxide dismutase SOD2 and different catalases and peroxidases were detected primarily in the dominant MAGs (Figure 2). No membrane-bound O_2_-reducing terminal oxidases (Cta, Cco, Cyo, Cyd) were detected (Supplementary Section 5).

## Discussion

Two planktonic, nitrification-inhibited denitrifying communities co-respiring O_2_ and nitrogen oxides were enriched under alternating oxic/anoxic conditions at frequencies representative of both natural (e.g. coastal sediments^20^) and engineered (e.g. wastewater treatment; Supplementary Section 6) ecosystems. Significant denitrification occurred at high oxygen concentrations, with almost 40% of the electrons from organic carbon being respired with NO_3_^-^ in the reactor with longer oxic/anoxic periods (R_4_). The high aerobic NO_3_^-^ reduction rates in this reactor -only half of the anoxic rates - suggest the enrichment of a more O_2_-tolerant denitrifying community than under more frequent oxic/anoxic transitions (R_32_). Typically, aerobic denitrification is characterized in monocultures under continuous aeration, resulting in relatively low reported rates (reviewed by Chen & Strous^8^). Only Patureau and colleagues have emphasized the significance of alternating oxic/anoxic conditions for enhanced aerobic denitrification^21^. However, all studies are based on a limited number of isolates, making their extrapolation to complex communities challenging. Few works quantified the contribution of aerobic denitrification in natural ecosystems with fluctuating oxic/anoxic conditions, namely aggregate-forming extracted soil bacteria^26^, sea sediments^25^, and coastal sediments^20^, yet at usually lower oxygen concentrations. Interestingly, Marchant et al.^20^ reported peaks of aerobic NO_3_^-^ reduction rates up to 60% of the anoxic rates at alternating oxic/anoxic conditions above 3 mg O_2_/L. However, only up to 5% of the electrons were respired via denitrification during aeration^20^, and anoxic niches could not be completely ruled out in any of the abovementioned studies. Overall, to the best of our knowledge, our results quantitatively prove for the first time that aerobic denitrification is an ecologically relevant process in microbial communities exposed to O_2_ fluctuations. Furthermore, we estimated that on average 11% (R_4_) and 22% (R_32_) of NO_3_^-^ was emitted as N_2_O during aeration, highlighting that aerobic denitrification also holds the potential to be a major contributor to aerobic N_2_O emissions.

The aerobic and anoxic proteomic profiles were nearly identical within each enrichment. The three most abundant MAGs in R_4_ and R_32_ accounted for 90% and 68% of the respective proteomes, proving their prominent functional role. All denitrification enzymes remained present and, at least partially, active under aerobic conditions. In contrast, in continuous monocultures, most denitrifying proteins are generally detected exclusively in anoxically-grown cells, and their abundance and activity is negligible under solely aerobic conditions^7,27,28^. Oxygen is known to suppress the transcription of denitrifying genes^7,24^, even if denitrification transcripts have also been detected during aeration^17,20,24,29^, and prolonged exposure to alternating conditions has been hypothesized to reduce the impact of O_218,20,21,26_. The constitutive detection of denitrifying enzymes is likely the direct result of the oxic/anoxic transition frequencies being significantly higher than the imposed growth rates, irrespective of potential transcriptional regulation mechanisms. In analogy, relevant aerobic residual denitrification potentials are to be expected in environments with rapid O_2_ fluctuations, such as sediments^20^ and wastewater treatment plants (Supplementary Section 6). The lower aerobic denitrification rates, compared to the anoxic ones, can thus reasonably be ascribed to reversible enzyme inhibition or electron competition with O_2_, rather than to transcriptional or translational regulation^8,30,31^. The O_2_ impact differed for each denitrification step, in line with previous observations^7,16^. While NO_2_^-^ and NO were hardly detected, N_2_O consistently accumulated, confirming the higher relative oxygen sensitivity of NosZ^16,26,32^. The marked N_2_O accumulation at the onset of anoxia implies a slower post-aerobiosis recovery of Nos compared to the other reductases. The progressive N_2_O accumulation under full aeration (Supplementary Figure S2) suggests a gradual yet incomplete inhibition of N_2_O reduction, as observed also by Qu et al.^31^. In fact, we estimated that 80-90% of the produced N_2_O was still reduced during aeration. Taken together, these results highlight the need for more research on the impact of fluctuating O_2_ on denitrification and, from a physiological perspective, further highlight the long-term competitive advantage of metabolic preparedness in dynamic environments.

Contrary to the long-standing assumption that the periplasmic reductase Nap is required for aerobic denitrification^20,21,23,26,33^, only the membrane-bound Nar was detected in our metaproteomes. Although preferential extraction or sequencing, and biases towards more abundant species can impact protein recovery^34^, both Nap subunits are soluble^35^ and are usually detected with equivalent protocols (e.g.: in *Paracoccus denitrificans*^28^). Also, the *napAB* genes were found in the most abundant MAGs, *e*.*g*. bin1.1 accounting for 50% of the proteome in R_4_. Therefore, while the presence of Nap at very low abundance cannot be completely ruled out, NO_3_^-^ reduction in our cultures was evidently driven by Nar and thus contributed directly to proton translocation under aerobic conditions. Studies on pure cultures of *Paracoccus pantotrophus* and *P. denitrificans* reported Nar and Nap to be preferentially expressed under continuous anoxic or aerobic conditions, respectively^28,33,36^. The excess NO_3_^-^ in our cultures might have alleviated the potential oxygen inhibition of NO_3_^-^ uptake^37,38^, favouring the lower-affinity Nar over Nap^39^. However, Ellington et al.^40^ measured high levels of *nap* transcription and Nap activity in *P. pantotrophus* grown in aerobic NO_3_^-^-excess chemostats, suggesting that factors other than NO_3_^-^ affinity determined the preferential Nar expression in our enrichments. Overall, the here observed consistent and exclusive expression of Nar suggests a higher versatility under alternating oxic/anoxic conditions, and challenges the use of *nap* as specific marker gene for aerobic denitrification^19,20^.

The subsequent nitrogen oxides reduction steps featured different degrees of labour-division among the MAGs in the two enrichments. Both nitrite reductases (NirK and NirS), and both clade I and II N_2_O reductase (NosZ) were primarily expressed by the dominant MAGs in R_4_. Conversely, the proteomic profile of R_32_ revealed a more prominent role of lower abundant MAGs in NO_2_^-^ and N_2_O reduction. Also, despite the widespread presence of the *nirK* gene in R_32_, mainly NirS was expressed. While O_2_-driven preferential expression of either NirK or NirS is plausible, conflicting O_2_-sensitivities have been reported^16^, warranting further research on the determinants of functional homologues preferences. In line with previous proteomic studies^28,41^, the detection of the membrane-bound hydrophobic qNor and cNor, intrinsically challenging to detect in proteomic analyses, was negligible. Interestingly, the *nosZ I* was annotated in most MAGs, with many expressing the encoded NosZ I. In turn, NosZ II was exclusively detected in R_4_. It is here tempting to speculate that the higher aerobic denitrification rates in R_4_ related to the reported lower O_2_ inhibition of clade II NosZ^17^. However, the observations from Suenaga et al. were limited to one *nosZ II*-harbouring *Azospira* strain and no evident clade-dependent differences in O_2_-tolerance were observed by Wang et al.^42^.

Furthermore, different physiological mechanisms such as strain-specific ability to scavenge O_2_ may impact the O_2_-tolerance of N_2_O-reducers^42^. In conclusion, these results reveal a much broader and unexplored biochemical breadth of aerobic denitrifiers in mixed communities, and quantitatively prove that aerobic denitrification cannot be neglected anymore in complex microbiomes with time-varying oxygen availabilities. We also posit that nitrifiers contribution to aerobic N_2_O emissions in natural and engineered ecosystems might currently be overestimated.

## Supporting information

Full Supplementary Information

## Acknowledgements

The authors are highly indebted to Dimitry Sorokin and Gijs Kuenen (TU Delft) for inspiring discussions and valuable feedback, Dirk Geerts and Dita Heikens (TU Delft) for precious support with the bioreactors and preparation of proteomic samples, Francesc Corbera Rubio (TU Delft) for providing the nitrifying culture, Waternet for providing the activated sludge, and Alexandra Deeke (Waterschap de Dommel), Cora Uijterlinde (STOWA), Inge Pistorius and Robert Kras (Waterschap Aa en Maas), Maaike Hoekstra (HHNK), Marcel Zandvoort (Waternet), and Mariska Ronteltap (Hoogheemraadschap van Delfland) for insightful discussions. The work was financed by Stichting Toegepast Onderzoek Waterbeheer (STOWA; JG191217009/732.750/CU), Hoogheemraadschap Hollands Noorderkwartier (HHNK; 20.0787440) and Waterschap de Dommel (Z62737/U131154). ML was supported by a Veni grant from the Dutch Research Council (NWO; project number VI.Veni.192.252).

## Materials and methods

### 1. Continuous-flow stirred tank reactors operation

Two 1 L jacketed continuous-flow stirred tank reactors (Applikon, Getinge) were operated during 96 days, with continuous mixing at 500 rpm using a six-blade turbine. The hydraulic and sludge retention times (HRT and SRT) were identical, controlled at 2 ± 0.1 days by two peristaltic pumps (Masterflex) continuously feeding the two media to the system and an effluent pump removing 94 mL of broth every 6 h. The average working volume was 0.75 ± 0.05 L. The temperature was controlled at 20 ± 0.1 °C using a cryostat bath (Lauda). The pH and dissolved oxygen were continuously monitored by pH and dissolved oxygen probes (Applikon AppliSens, Getinge). The pH was kept at 7.1 ± 0.1 by 1 M HCl or 1 M NaOH with two peristaltic pumps (Watson Marlow) controlled by a process controller (Applikon in-Control, Getinge).

Denitrifying bacteria were enriched by continuous supply of 0.93 ± 0.04 N-mmol/h NO_3_^-^ as electron acceptor and a mixture of volatile fatty acids (VFAs) as electron donor and carbon source: acetate (0.94 ± 0.08 C-mmol/h), propionate (1.00 ± 0.09 C-mmol/h) and butyrate (0.75 ± 0.07 C-mmol/h). After the initial start-up phase of 27 days, the VFAs were always below detection limit in the effluent, confirming carbon limiting conditions. Ammonia served as the nitrogen source. The reactors were covered with aluminium foil to prevent the growth of phototrophic organisms. Nitrogen and carbon media were prepared separately to prevent microbial growth during storage. Nitrogen medium consisted of (per liter): 9.14 g NaNO_3_, 2.84 g NH_4_Cl, 2.01 g KH_2_PO_4_, 1.04 g MgSO_4_ · 7 H_2_O, 0.04 g NaOH, 4 mg yeast extract, 5 mL trace element solution^43^ and 1 mL of a 10 g/L solution of allylthiourea (ATU). ATU selectively inhibits bacterial ammonium oxidation to nitrite^44–46^ without significantly affecting denitrification^47^. The trace element solution consisted of (per liter): 50 g EDTA · H_2_ · Na_2_ · 2 H_2_O, 2.5 g FeSO_4_ · 7 H_2_O, 1.1 g ZnSO_4_ · 7 H_2_O, 4.1 g CaCl_2_ · 2 H_2_O, 2.2 g MnSO_4_ · H_2_O, 1.1 g Na_2_MoO_4_ · 2 H_2_O, 0.8 g CuSO_4_ · 5 H_2_O and 0.7 g CoCl_2_ · 6 H_2_O. Carbon medium consisted of (per liter): 8.1 g NaCH_3_OO · 3 H_2_O, 1.9 mL C_4_H_8_O_2_ and 4.1 g NaC_3_H_5_O_2,_ the pH was set to 6.0 with NaOH pellets. Drops of antifoam C emulsion (Merck Life Science NV), diluted six times, was added to the reactors when foam formation was noted.

The reactors were inoculated with activated sludge from the Amsterdam-West wastewater treatment plant and run under NO_3_^-^-limiting conditions for 20 days before switching to carbon-limiting conditions. During the NO_3_^-^-limiting start-up phase, the concentration of VFAs were increased by four times in the carbon medium compared to the values presented above. The two reactors were exposed to continuous cycles of alternating aerobic and anoxic conditions in a time proportion of 2:1. The reactors were exposed to 4 (R_4_) or 32 (R_32_) cycles per day, with aerobic periods of 4h and 30 min and anoxic periods of 2h and 15 min, respectively. Aerobic and anoxic conditions were maintained by continuous sparging of compressed air and N_2_, respectively, at 400 mL/min, controlled by mass-flow controllers (Brooks). Aerobic conditions close to air saturation were assured by maintaining average dissolved O_2_ concentrations of 7.5 ± 0.2 and 6.8 ± 0.3 mg/L in R_4_ and R_32_, respectively. The reactors reached fully anoxic or aerobic conditions within 5 min after switching the influent gas. The 6-hourly reactor broth removal coincided with the end of an anoxic phase. Considering the transition periods, the net amount of aerobic (>= 1% air saturation) and anoxic (< 1% air saturation) hours per day were 16:8 and 17:7 for R_4_ and R_32_, respectively. Throughout the steady-state operation of R_4_ and R_32_, visual analysis confirmed that the cultures remained suspended and well-mixed. For R_32_, small biomass aggregates were progressively washed out reaching an entirely homogeneous and suspended culture after 63 days of operation. Occasional wall growth was regularly removed, with no noticeable impact on reactor performance (Supplementary Figure S1). For metabolite and biomass analysis, quadruplicate samples of 2 mL were taken from both reactors at three moments within a cycle: at the start and end of the aerobic phase and at the end of the anoxic phase. The samples were placed on ice and immediately filtered using 0.22 μm PVDF Millex-GV syringe filters (Merck) or centrifuged at 16,200 x *g* for 5 minutes at 4 °C to separate the biomass from the supernatant. The pellets were stored at -80 °C and the supernatant at -20 °C until further analysis. Feed substrate concentrations were confirmed by occasionally sampling the reactor influent, with storage at -20 °C until analysis.

### 2. Analytical methods

The concentrations of NH_4_^+^, NO_2_^-^ and NO_3_^-^ in the influent and effluent supernatant were spectrophotometrically measured with the Gallery™ Discrete Analyzer (Thermo Fisher Scientific) or cuvette test kits (Hach Lange) immediately after sampling or within 24h after storage at 4 °C. The concentrations of acetate, propionate and butyrate in the influent and effluent supernatant were measured after storage at -20 °C by high-pressure liquid chromatography (Vanquish Core HPLC, Thermo Fisher Scientific) using an Aminex HPX-87H column (300 × 7.8 mm) (Bio-Rad), calibrated with solutions ranging from 0 to 250 mM. The concentrations of O_2_, N_2_O and CO_2_ in the off-gas were continuously monitored online by a Rosemount NGA 2000 off-gas analyser (Emerson). Before reaching the analyser, the off-gas was dried in a condenser, operated with water at 4 °C using a cryostat bath (Lauda).

### 3. Calculations

The calculations of consumption and production rates of all compounds are detailed in Supplementary Section 2. Briefly, the overall (*i*.*e*. the weighted average of oxic and anoxic) consumption and production rates of dissolved compounds (NH_4_^+^, NO_2_^-^, NO_3_^-^, acetate, propionate, and butyrate) were calculated via a mass balance of the volumetric influent and effluent flow rates, and the influent and effluent concentrations. The biomass production rate was estimated from the ammonium consumption rates, assuming complete assimilation into biomass at a ratio of 0.2 Nmol/Cmol. The same estimation was obtained when calculating the biomass rates from the carbon balance (*i*.*e*. from the CO_2_ and organic carbon rates), validating the previous assumption. The effluent concentrations were taken as the average of the three effluent measurements (beginning and end of the oxic phase and end of the anoxic phase). The overall, aerobic and anoxic accumulation rates of gaseous compounds (N_2_O and CO_2_) were calculated from continuous measurements (every minute) of the molar fractions in the off-gas. The aerobic phase was assumed to be the period in which the dissolved oxygen was above 1% air saturation. The overall N_2_ production rate was estimated from the nitrate and N_2_O rates, as the accumulation of nitrite and nitric oxide was negligible throughout steady-state. The O_2_ consumption rates during the aerobic phase were calculated from the continuous dissolved oxygen measurements and experimentally determined volumetric mass transfer coefficient (Supplementary Section 1). For consistency, an “overall” consumption rate was also calculated for O_2_, by averaging its aerobic consumption over the entire cycle duration. For all compounds, steady-state rates were determined by averaging the rates measured during the entire steady-state period. Overall carbon and electron balances were calculated from the consumption and production rates of all substrates and products. The specific aerobic and anoxic rates of the dissolved nitrogen compounds were estimated from the available measurements as explained in Supplementary Section 2. Possible deviations in the estimated rates due to potential PHA accumulation were negligible (Supplementary Section 2).

### 4. DNA extraction, library preparation and sequencing

DNA was extracted from biomass samples taken at the end of the anoxic period after 68 days of operation using the DNeasy PowerSoil Pro Kit (Qiagen), according to the manufacturer’s instructions with the following exceptions. The pelleted biomass, stored at -80 °C, was resuspended in 800 μL of solution CD1 by vortexing before transferring to the PowerBead tube. Samples were homogenized by 4 x 40s bead-beating using the Beadbeater-24 (Biospec) alternated with 2 min incubation on ice. Tubes were gently inverted 10x instead of vortexing to avoid DNA shearing. Elution of the extracted DNA was performed with 50 μL solution C6. The DNA concentration was 710 and 605 ng/μL for R_4_ and R_32_, respectively, as measured with the Qubit 4 Fluorometer (Thermo Fisher Scientific). DNA quality was assessed with the BioTek Synergy HTX multi-mode microplate reader (Agilent). For differential coverage binning and increased bin recovery, DNA was also extracted from samples taken after 41 days of operation using the DNeasy UltraClean Microbial Kit (Qiagen), following the manufacturer’s instructions. The extraction yielded 224 and 267 ng/μL for R_4_ and R_32_, respectively.

Library preparation of the extracted DNA from day 68 for long-read sequencing was performed using the Ligation Sequencing Kit V14 (Oxford Nanopore Technologies Ltd). The NEBNext^®^ Companion Module for Oxford Nanopore Technologies^®^ Ligation Sequencing (New England BioLabs Inc.) and UltraPure™ BSA (50 mg/mL) (Thermo Fisher Scientific) were additionally used for the DNA repair and end-prep and the flow cell priming steps. All steps were performed as instructed by the manufacturer, except the incubations in the Hula mixer were replaced with slow manual inversions (∼5 s per inversion). All resuspension steps were performed by flicking the tube. MinION R10.4 version flow cells (Oxford Nanopore), starting with 1345 and 461 active pores, were loaded with 132 and 150 ng DNA for R_4_ and R_32_, respectively. Samples were sequenced in accurate mode (260 bps) for 46 and 40 h, respectively, yielding 14.7 and 4.3 Gbp of sequenced data. Samples from day 41 were sequenced on a Illumina NovaSeq 6000 platform by Novogene Ltd. (UK). Approximately 10 Gbp of 150 bp paired-end reads with an insert size of 350 bp were generated.

### 5. Metagenomic data processing

The raw Nanopore data was basecalled using Guppy v6.4.2 (Oxford Nanopore) with the configuration file “dna_r10.4.1_e8.2_260bps_sup.cfg” and --do_read_splitting option. Duplex reads were identified and filtered using the pairs_from_summary and filter_pairs settings from Duplex tools v0.2.19 (Oxford Nanopore) and basecalled with the duplex basecaller of Guppy, using identical settings to the simplex basecalling. The simplex reads, not part of a pair, were merged with the duplex basecalled reads using SeqKit v2.3.0^48^, generating a single fastq file containing all unique reads. Sequences belonging to the Lambda control DNA were removed with NanoLyse v1.2.1^49^. The basecalled data was inspected with NanoPlot v1.41.0^49^. Reads were filtered with NanoFilt -q 10 -l 1000 (v2.8.0^49^) and trimmed with Porechop v0.2.4 (https://github.com/rrwick/Porechop). Reads assembly was performed with Flye v2.9.1 in -- meta mode. Assembly quality was assessed with MetaQUAST v5.0.2^50^ using the --fragmented option. Reads were aligned to the assembly with Minimap2 v2.24^51^. The assembly was polished with Racon v1.4.3 (https://github.com/isovic/racon) and two rounds of Medaka v1.5.0 (https://github.com/nanoporetech/medaka) with default settings. Nanopore and Illumina reads were mapped to the final assembly using Minimap2, the alignments were converted from SAM to BAM and sorted with SAMtools v1.10^52^, and the contig coverage was calculated with jgi_summarize_bam_contig_depths^53^. Automatic differential coverage binning was independently performed with MetaBAT2 v2.15^53^, MaxBin2 v2.2.7^54^ and CONCOCT v1.1.0^55^, with a minimum contig length of 2000 bp. The output of all binning tools was combined with DAS Tool v1.1.3^56^, using Prodigal and DIAMOND v2.0.8^57^ for single copy gene prediction and identification, resulting in an optimized non-redundant set of bins. Bin completeness and contamination was determined with CheckM v1.1.3^58^ using the lineage_wf workflow. Nanopore and Illumina bins from each reactor were dereplicated with dRep v3.2.2^59^ with the options -comp 70 -con 10 --S_algorithm gANI, using the default thresholds for average nucleotide identity (ANI). The final set of non-redundant bins (completeness above 70% and contamination under 10%) contained all Nanopore bins and the Illumina bins that did not cluster with any Nanopore bins (gANI < 99%). The bins were taxonomically classified with the classify_wf workflow of GTDB-Tk v.2.2.5^60^ using the GTDB release 207 (gtdbtk_r207_v2_data.tar.gz^61^). The relative abundance of each bin in the metagenome was determined with CoverM v0.6.1 (https://github.com/wwood/CoverM) in relative_abundance mode.

Genes were predicted from the assembly using Prodigal v2.6.3^62^ and functionally annotated with DRAM in annotate_genes mode^63^, using the default settings and the KOfam^64^, MEROPS^65^, Pfam^66^, dbCAN^67^, and VOGDB (https://vogdb.org/) databases. Genes of interest were identified by their KO identifier (Supplementary Tables S7, S8 and S9). The genes encoding the alpha and beta subunits of the respiratory nitrate reductase (Nar) have the same KO identifiers as the alpha and beta subunits of the nitrite oxidoreductase (Nxr). We could confidently attribute all genes identified with K00370 and K00371 to the nitrate reductase (encoded by *narGHI* or *narZYV*), as the gamma subunit of this enzyme (K00374, exclusive to Nar) was present in all bins containing the alpha and beta subunits. Distinction between clade I and clade II N_2_O reductase (NosZ) was determined by, respectively, identifying the twin-arginine translocation (Tat, IPR006311) or the general secretory (Sec, IPR026468) pathway-specific signal peptides on InterPro^68^. The quinol-dependent nitric oxide reductase (qNor, encoded by *norZ*) has a fused quinol oxidase domain on the N-terminal^69^, unlike the cytochrome c-dependent reductase (cNor, encoded by *norBC*). Yet, the *norZ* genes were annotated as *norB*, so qNor was distinguished by identifying the quinol oxidase domain through a multiple sequence alignment of putative NorB protein sequences (K04561) with reference sequences of NorB (*Pseudomonas stutzeri*, P98008) and NorZ (*Cupriavidus necator*, Q0JYR9), extracted from UniProtKB^70^.

Quality control of the Illumina paired-end reads was performed with FastQC v0.11.7 (https://www.bioinformatics.babraham.ac.uk/projects/fastqc/). Reads were filtered and trimmed with Trimmomatic v0.39^71^ using the options LEADING:3 TRAILING:3 SLIDINGWINDOW:4:15 MINLEN:35 HEADCROP:5. Reads were assembled into contigs using metaspades.py from SPAdes v3.14.1^72^. The assembly was inspected with MetaQUAST v5.0.2 using the --fragmented option. Contigs smaller than 500 bp were removed with filterContigByLength.pl^73^. Gene prediction and functional annotation was performed identically to the Nanopore data. The paired-end reads were mapped to the contigs using BWA-MEM2 v2.1^74^ and the alignments were processed as described above. Automatic binning and bin analysis was identical as described for the Nanopore data, except no differential coverage binning was performed and the default minimum contig length of each binning software was used. The generated bins were further dereplicated with the Nanopore bins as described above. Nonpareil v3.401^75^, ran with the kmer algorithm, estimated that the Illumina reads covered 98.6% and 99.2% of the sample diversity.

### 6. Protein extraction, precipitation, digestion and clean-up

Preparation of protein samples was performed according to Kleikamp et al.^76^. Briefly, biomass samples were homogenised with glass beads (150 - 212 μm, Sigma Aldrich), 50 mM TEAB buffer with 1% (w/w) NaDOC and B-PER reagent (Thermo Scientific) through three cycles of vortexing and ice incubation. The samples were incubated at 80 °C and sonicated. The supernatant was collected after centrifuging at 14,000 g. Proteins were precipitated with 1:4 trichloroacetic acid solution (TCA, Sigma Aldrich) and washed with acetone. The pellet was re-dissolved in 6 M Urea (Sigma Aldrich) in 200 mM ammonium bicarbonate, reduced in 10 mM dithiothreitol (Sigma Aldrich) at 37 °C for 60 min, and alkylated with 20 mM iodoacetamide (Sigma Aldrich) in the dark for 30 min, at room temperature. Samples were diluted to reach a urea concentration under 1 M. Proteins were digested overnight (21 h) at 37 °C with 0.1 μg/μL trypsin (sequencing grade, Promega) dissolved in 1 mM HCl. Samples were desalted and cleaned through solid phase extraction using an Oasis HLB 96-well μElution Plate (2mg sorbent per well, 30 μm, Waters) and a vacuum pump. The columns were conditioned with MeOH, equilibrated with two rounds of water, loaded with the digested samples and washed with two rounds of 5% MeOH. Peptide samples were sequentially eluted with 2% formic acid in 80% MeOH and 1 mM ammonium bicarbonate in 80% MeOH, dried at 50 °C in an Integrated SpeedVac™ System (Thermo Scientific) and stored at -20 °C until shotgun proteomic analysis.

### 7. Shotgun metaproteomics

Briefly, samples were dissolved in 20 μL of 3% acetonitrile and 0.01% trifluoroacetic acid. The samples were incubated at room temperature for 30 min and vortexed thoroughly. The protein concentration was measured on a NanoDrop ND-1000 spectrophotometer (Thermo Scientific) at 280 nm wavelength. If needed, samples were diluted to a concentration of 0.5 mg/mL.

Shotgun metaproteomics experiments were performed as recently described^76,77^. Briefly, aliquots corresponding to approximately 0.5 μg protein digest were analysed using a nano-liquid-chromatography system consisting of an EASY nano-LC 1200, equipped with an Acclaim PepMap RSLC RP C18 separation column (50 μm x 150 mm, 2 μm, Cat. No. 164568), and a QE plus Orbitrap mass spectrometer (Thermo Fisher Scientific). The flow rate was maintained at 350 nL/min over a linear gradient from 5% to 25% solvent B over 90 min, then from 25% to 55% over 60 min, followed by back equilibration to starting conditions. Solvent A was H_2_O containing 0.1% formic acid (FA), and solvent B consisted of 80% ACN in H_2_O and 0.1% FA. The Orbitrap was operated in data dependent acquisition (DDA) mode acquiring peptide signals from 385–1250 m/z at 70 K resolution in full MS mode with a maximum ion injection time (IT) of 75 ms and an automatic gain control (AGC) target of 3E6. The top 10 precursors were selected for MS/MS analysis and subjected to fragmentation using higher-energy collisional dissociation (HCD) at a normalised collision energy of 28. MS/MS scans were acquired at 17.5 K resolution with AGC target of 2E5 and IT of 75 ms, 1.2 m/z isolation width. Raw mass spectrometric data from each reactor were analysed against a protein reference sequence database respectively constructed from the metagenomic data, including the all MAGs and unbinned portion of the samples taken at day 68 and the additional dereplicated MAGs from day 41, using PEAKS Studio X (Bioinformatics Solutions Inc.) allowing for 20 ppm parent ion and 0.02 m/z fragment ion mass error, 3 missed cleavages, and iodoacetamide as fixed and methionine oxidation and N/Q deamidation as variable modifications. Peptide spectrum matches were filtered against 1% false discovery rates (FDR) and protein identifications with ≥2 unique peptide sequences.

For each protein, the peptide spectral counts were normalized by dividing them with the protein molecular weight. The relative abundance of each protein in the samples was calculated by dividing its normalized spectral counts by the sum of normalized spectral counts of all proteins of that respective sample. The technical duplicates were then averaged. The total relative contribution of each bin to the proteome was determined by summing the relative abundances of its proteins. Similarly, the total relative abundance of functionally identical proteins was determined by summing the relative contribution of all proteins with the same functional annotation. The exclusion of any NapA and NapB peptides in the proteomic data was concluded from the absence of corresponding sequences within the obtained peptide spectrum matches. RStudio v22.0.3^78^ with R v4.2.2^79^, with the plyr^80^, tidyverse^81^, readxl^82^ and ggplot2^83^ packages, was used for data processing and visualization.

## References

1. IPCC. Climate Change 2014: Synthesis Report. (2014).

2. Tian, H. et al. A comprehensive quantification of global nitrous oxide sources and sinks. Nature 586, 248–256 (2020).

3. Freing, A., Wallace, D. W. R. & Bange, H. W. Global oceanic production of nitrous oxide. Philos. Trans. R. Soc. B 367, 1245–1255 (2012).

4. Zhu, X., Burger, M., Doane, T. A. & Horwath, W. R. Ammonia oxidation pathways and nitrifier denitrification are significant sources of N2O and NO under low oxygen availability. Proc. Natl. Acad. Sci. U. S. A. 110, 6328–6333 (2013).

5. Duan, H. et al. Mitigating nitrous oxide emissions at a full-scale wastewater treatment plant. Water Res. 185, 116196 (2020).

6. Duan, H. et al. Insights into Nitrous Oxide Mitigation Strategies in Wastewater Treatment and Challenges for Wider Implementation. Environ. Sci. Technol. (2021) doi:10.1021/acs.est.1c00840.

7. Korner, H. & Zumft, W. G. Expression of denitrification enzymes in response to the dissolved oxygen levels and respiratory substrate in continuous culture of Pseudomonas stutzeri. Appl. Environ. Microbiol. 55, 1670–1676 (1989).

8. Chen, J. & Strous, M. Denitrification and aerobic respiration, hybrid electron transport chains and co-evolution. Biochim. Biophys. Acta - Bioenerg. 1827, 136–144 (2013).

9. Braker, G. & Conrad, R. Diversity, structure, and size of N 2 O-producing microbial communities in soils-what matters for their functioning? Advances in Applied Microbiology vol. 75 (2011).

10. Baggs, E. M. Soil microbial sources of nitrous oxide: Recent advances in knowledge, emerging challenges and future direction. Curr. Opin. Environ. Sustain. 3, 321–327 (2011).

11. Butterbach-Bahl, K., Baggs, E. M., Dannenmann, M., Kiese, R. & Zechmeister-Boltenstern, S. Nitrous oxide emissions from soils: How well do we understand the processes and their controls? Philos. Trans. R. Soc. B Biol. Sci. 368, (2013).

12. Battaglia, G. & Joos, F. Marine N2O Emissions From Nitrification and Denitrification Constrained by Modern Observations and Projected in Multimillennial Global Warming Simulations. Global Biogeochem. Cycles 32, 92–121 (2018).

13. Kampschreur, M. J., Temmink, H., Kleerebezem, R., Jetten, M. S. M. & van Loosdrecht, M. C. M. Nitrous oxide emission during wastewater treatment. Water Res. 43, 4093–4103 (2009).

14. van Loosdrecht, M. C. M. & Jetten, M. S. M. Microbiological conversions in nitrogen removal. Water Sci. Technol. 38, 1–7 (1998).

15. Gruber, W. et al. Tracing N2O formation in full-scale wastewater treatment with natural abundance isotopes indicates control by organic substrate and process settings. Water Res. X 15, (2022).

16. Otte, S., Grobben, N. G., Robertson, L. A., Jetten, M. S. M. & Kuenen, J. G. Nitrous oxide production by Alcaligenes faecalis under transient and dynamic aerobic and anaerobic conditions. Appl. Environ. Microbiol. 62, 2421–2426 (1996).

17. Suenaga, T., Riya, S., Hosomi, M. & Terada, A. Biokinetic characterization and activities of N2O-reducing bacteria in response to various oxygen levels. Front. Microbiol. 9, 1–10 (2018).

18. Robertson, L. A. & Kuenen, J. G. Aerobic denitrification: a controversy revived. Arch. Microbiol. 139, 351–354 (1984).

19. Yang, J. et al. A critical review of aerobic denitrification: Insights into the intracellular electron transfer. Sci. Total Environ. 731, 139080 (2020).

20. Marchant, H. K. et al. Denitrifying community in coastal sediments performs aerobic and anaerobic respiration simultaneously. ISME J. 11, 1799–1812 (2017).

21. Patureau, D., Zumstein, E., Delgenes, J. P. & Moletta, R. Aerobic denitrifiers isolated from diverse natural and managed ecosystems. Microb. Ecol. 39, 145–152 (2000).

22. Frette, L., Gejlsbjerg, B. & Westermann, P. Aerobic denitrifiers isolated from an alternating activated sludge system. FEMS Microbiol. Ecol. 24, 363–370 (1997).

23. Carter, J. P., Ya Hsin Hsiao, Spiro, S. & Richardson, D. J. Soil and sediment bacteria capable of aerobic nitrate respiration. Appl. Environ. Microbiol. 61, 2852–2858 (1995).

24. Giannopoulos, G. et al. Tuning the modular Paracoccus denitrificans respirome to adapt from aerobic respiration to anaerobic denitrification. Environ. Microbiol. 19, 4953–4964 (2017).

25. Gao, H. et al. Aerobic denitrification in permeable Wadden Sea sediments. ISME J. 4, 417–426 (2010).

26. Morley, N., Baggs, E. M., Dörsch, P. & Bakken, L. Production of NO, N2O and N2 by extracted soil bacteria, regulation by NO2- and O2 concentrations. FEMS Microbiol. Ecol. 65, 102–112 (2008).

27. Moir, J. W. B., Richardson, D. J. & Ferguson, S. J. The expression of redox proteins of denitrification in Thiosphaera pantotropha grown with oxygen, nitrate, and nitrous oxide as electron acceptors. Arch. Microbiol. 164, 43–49 (1995).

28. Olaya-Abril, A. et al. Exploring the denitrification proteome of Paracoccus denitrificans PD1222. Front. Microbiol. 9, 1–16 (2018).

29. Abada, A. et al. Aerobic bacteria produce nitric oxide via denitrification and promote algal population collapse. ISME J. 1–17 (2023) doi:10.1038/s41396-023-01427-8.

30. Conthe, M., Parchen, C., Stouten, G., Kleerebezem, R. & van Loosdrecht, M. C. M. O2 versus N2O respiration in a continuous microbial enrichment. Appl. Microbiol. Biotechnol. 102, 8943–8950 (2018).

31. Qu, Z., Bakken, L. R., Molstad, L., FrostegÅrd, Å. & Bergaust, L. L. Transcriptional and metabolic regulation of denitrification in Paracoccus denitrificans allows low but significant activity of nitrous oxide reductase under oxic conditions. Environ. Microbiol. 18, 2951–2963 (2016).

32. Wüst, A. et al. Nature’s way of handling a greenhouse gas: The copper-sulfur cluster of purple nitrous oxide reductase. Biol. Chem. 393, 1067–1077 (2012).

33. Bell, L. C., Richardson, D. J. & Ferguson, S. J. Periplasmic and membrane-bound respiratory nitrate reductases in Thiosphaera pantotropha. FEBS 265, 85–87 (1990).

34. Kleiner, M. et al. Assessing species biomass contributions in microbial communities via metaproteomics. Nat. Commun. 8, (2017).

35. Arnoux, P. et al. Structural and redox plasticity in the heterodimeric periplasmic nitrate reductase. Nat. Struct. Biol. 10, 928–934 (2003).

36. Richardson, D. J., Berks, B. C., Russell, D. A., Spiro, S. & Taylor, C. J. Functional, biochemical and genetic diversity of prokaryotic nitrate reductases. Cell. Mol. Life Sci. 58, 165–178 (2001).

37. Hernandez, D. & Rowe, J. J. Oxygen inhibition of nitrate uptake is a general regulatory mechanism in nitrate respiration. J. Biol. Chem. 263, 7937–7939 (1988).

38. Moir, J. W. B. & Wood, N. J. Nitrate and nitrite transport in bacteria. Cell. Mol. Life Sci. 58, 215–224 (2001).

39. Potter, L. C., Millington, P., Griffiths, L., Thomas, G. H. & Cole, J. A. Competition between Escherichia coli strains expressing either a periplasmic or a membrane-bound nitrate reductase: Does Nap confer a selective advantage during nitrate-limited growth? Biochem. J. 344, 77–84 (1999).

40. Ellington, M. J. K. et al. Characterization of the expression and activity of the periplasmic nitrate reductase of Paracoccus pantotrophus in chemostat cultures. Microbiology 149, 1533–1540 (2003).

41. Zorz, J. K., Kozlowski, J. A., Stein, L. Y., Strous, M. & Kleiner, M. Comparative proteomics of three species of ammonia-oxidizing bacteria. Front. Microbiol. 9, 1–15 (2018).

42. Wang, Z., Vishwanathan, N., Kowaliczko, S. & Ishii, S. Clarifying Microbial Nitrous Oxide Reduction Under Aerobic Conditions : Tolerant, Intolerant, and Sensitive. Microbiol. Spectr. (2023).

43. Vishniac, W. & Santer, M. The Thiobacilli. Bacteriol. Rev. (1957).

44. van Loosdrecht, M. C. M., Nielsen, P. H., Lopez-Vazquez, C. M. & Brdjanovic, D. Experimental Methods in Wastewater Treatment. Water Intelligence Online vol. 15 (2016).

45. Hooper, A. B. & Terry, K. R. Photoinactivation of ammonia oxidation in Nitrosomonas. J. Bacteriol. 119, 899–906 (1974).

46. Ginestet, P., Audic, J. M., Urbain, V. & Block, J. C. Estimation of nitrifying bacterial activities by measuring oxygen uptake in the presence of the metabolic inhibitors allylthiourea and azide. Appl. Environ. Microbiol. 64, 2266–2268 (1998).

47. Jensen, M. M., Thamdrup, B. & Dalsgaard, T. Effects of specific inhibitors on anammox and denitrification in marine sediments. Appl. Environ. Microbiol. 73, 3151–3158 (2007).

48. Shen, W., Le, S., Li, Y. & Hu, F. SeqKit: A cross-platform and ultrafast toolkit for FASTA/Q file manipulation. PLoS One 11, 1–10 (2016).

49. De Coster, W., D’Hert, S., Schultz, D. T., Cruts, M. & Van Broeckhoven, C. NanoPack: Visualizing and processing long-read sequencing data. Bioinformatics 34, 2666–2669 (2018).

50. Mikheenko, A., Saveliev, V. & Gurevich, A. MetaQUAST: Evaluation of metagenome assemblies. Bioinformatics 32, 1088–1090 (2016).

51. Li, H. Minimap2 : pairwise alignment for nucleotide sequences. 34, 3094–3100 (2018).

52. Li, H. et al. The Sequence Alignment/Map format and SAMtools. Bioinformatics 25, 2078–2079 (2009).

53. Kang, D. D. et al. MetaBAT 2: An adaptive binning algorithm for robust and efficient genome reconstruction from metagenome assemblies. PeerJ 2019, 1–13 (2019).

54. Wu, Y. W., Simmons, B. A. & Singer, S. W. MaxBin 2.0: An automated binning algorithm to recover genomes from multiple metagenomic datasets. Bioinformatics 32, 605–607 (2016).

55. Alneberg, J. et al. Binning metagenomic contigs by coverage and composition. Nat. Methods 11, 1144–1146 (2014).

56. Sieber, C. M. K. et al. Recovery of genomes from metagenomes via a dereplication, aggregation and scoring strategy. Nat. Microbiol. 3, 836–843 (2018).

57. Buchfink, B., Xie, C. & Huson, D. H. Fast and sensitive protein alignment using DIAMOND. Nat. Methods 12, 59–60 (2014).

58. Parks, D. H., Imelfort, M., Skennerton, C. T., Hugenholtz, P. & Tyson, G. W. CheckM: Assessing the quality of microbial genomes recovered from isolates, single cells, and metagenomes. Genome Res. 25, 1043–1055 (2015).

59. Olm, M. R., Brown, C. T., Brooks, B. & Banfield, J. F. DRep: A tool for fast and accurate genomic comparisons that enables improved genome recovery from metagenomes through de-replication. ISME J. 11, 2864–2868 (2017).

60. Chaumeil, P. A., Mussig, A. J., Hugenholtz, P. & Parks, D. H. GTDB-Tk v2: memory friendly classification with the genome taxonomy database. Bioinformatics 38, 5315–5316 (2022).

61. Parks, D. H. et al. GTDB: An ongoing census of bacterial and archaeal diversity through a phylogenetically consistent, rank normalized and complete genome-based taxonomy. Nucleic Acids Res. 50, D785–D794 (2022).

62. Hyatt, D. et al. Prodigal: Prokaryotic gene recognition and translation initiation site identification. BMC Bioinformatics 11, (2010).

63. Shaffer, M. et al. DRAM for distilling microbial metabolism to automate the curation of microbiome function. Nucleic Acids Res. 48, 8883–8900 (2020).

64. Aramaki, T. et al. KofamKOALA: KEGG Ortholog assignment based on profile HMM and adaptive score threshold. Bioinformatics 36, 2251–2252 (2020).

65. Rawlings, N. D., Waller, M., Barrett, A. J. & Bateman, A. MEROPS: The database of proteolytic enzymes, their substrates and inhibitors. Nucleic Acids Res. 42, 503–509 (2014).

66. Mistry, J. et al. Pfam: The protein families database in 2021. Nucleic Acids Res. 49, D412–D419 (2021).

67. Yin, Y. et al. DbCAN: A web resource for automated carbohydrate-active enzyme annotation. Nucleic Acids Res. 40, 445–451 (2012).

68. Paysan-Lafosse, T. et al. InterPro in 2022. Nucleic Acids Res. 51, D418–D427 (2023).

69. Zumft, W. G. Nitric oxide reductases of prokaryotes with emphasis on the respiratory, heme-copper oxidase type. J. Inorg. Biochem. 99, 194–215 (2005).

70. The UniProt Consortium. UniProt: the Universal Protein Knowledgebase in 2023. Nucleic Acids Res. 51, 523–531 (2023).

71. Bolger, A. M., Lohse, M. & Usadel, B. Trimmomatic: A flexible trimmer for Illumina sequence data. Bioinformatics 30, 2114–2120 (2014).

72. Nurk, S., Meleshko, D., Korobeynikov, A. & Pevzner, P. A. MetaSPAdes: A new versatile metagenomic assembler. Genome Res. 27, 824–834 (2017).

73. Dong, X. & Strous, M. An Integrated Pipeline for Annotation and Visualization of Metagenomic Contigs. Front. Genet. 10, 1–10 (2019).

74. Md, V., Misra, S., Li, H. & Aluru, S. Efficient Architecture-Aware Acceleration of BWA-MEM for Multicore Systems. IEEE Parallel Distrib. Process. Symp. (2019) doi:10.1109/IPDPS.2019.00041.

75. Rodriguez-R, L. M. & Konstantinidis, K. T. Nonpareil: A redundancy-based approach to assess the level of coverage in metagenomic datasets. Bioinformatics 30, 629–635 (2014).

76. Kleikamp, H. B. C. et al. Database-independent de novo metaproteomics of complex microbial communities. Cell Syst. 12, 375-383.e5 (2021).

77. den Ridder, van den Brandeler, W., Altiner, M., Daran-Lapujade, P. & Pabst, M. Proteome dynamics during transition from exponential to stationary phase under aerobic and anaerobic conditions in yeast. Mol. Cell. Proteomics 143747 (2023) doi:10.1016/j.mcpro.2023.100552.

78. RStudio Team. RStudio: Integrated Development Environment for R. RStudio, PBC, Boston, MA http://www.rstudio.com/ (2021).

79. R Core Team. R: A language and environment for statistical computing. R Foundation for Statistical Computing, Vienna, Austria https://www.r-project.org/ (2022).

80. Wickham, H. The split-apply-combine strategy for data analysis. J. Stat. Softw. 40, 1–29 (2011).

81. Wickham, H. et al. Welcome to the Tidyverse. J. Open Source Softw. 4, 1686 (2019).

82. Wickham, H. & Bryan, J. readxl: Read Excel Files. https://cran.r-project.org/package=readxl (2023).

83. Wickham, H. ggplot2: Elegant Graphics for Data Analysis. Springer-Verlag New York. Media vol. 35 (2016).

