## Supplementary material for "Aerobic denitrification as N_2_O source in microbial communities": Full Supplementary Information

### 1. Reactor operation

**Table S1. Measured average substrate loading and steady-state conversion rates of the low- ($R_4$) and high-frequency ($R_{32}$) reactors.** Overall rates refer to rates estimated over the total duration of an oxic/anoxic cycle, and considers the average of three effluent concentrations (beginning and end of aerobic phase, and end of anoxic one). Only for the gaseous compounds ($CO_2$ and $N_2O$) individual rates for each phase (aerobic and anoxic) were measured on top of the overall rates. $O_2$ was only added and consumed in the aerobic phase, yet an "overall" rate was also calculated by averaging the aerobic $O_2$ consumption over the entire cycle duration (eq. S9) for further balancing purposes.

| | | | Loading | | Conversion | |
|---|---|---|---|---|---|---|
| Compound | Phase | Units | $R_4$ | $R_{32}$ | $R_4$ | $R_{32}$ |
| $CO_2$ | Aerobic | C-mmol/h | - | - | 1.37 ± 0.07 | 1.5 ± 0.1 |
| | Anoxic | C-mmol/h | - | - | 1.15 ± 0.08 | 1.4 ± 0.1 |
| | Overall | C-mmol/h | - | - | 1.30 ± 0.06 | 1.5 ± 0.1 |
| $N_2O$ | Aerobic | N-mmol/h | - | - | 0.06 ± 0.04 | 0.04 ± 0.03 |
| | Anoxic | N-mmol/h | - | - | 0.04 ± 0.04 | 0.04 ± 0.02 |
| | Overall | N-mmol/h | - | - | 0.05 ± 0.03 | 0.04 ± 0.03 |
| $NO_3^-$ | Overall | N-mmol/h | 0.96 ± 0.02 | 0.91 ± 0.02 | -0.73 ± 0.08 | -0.60 ± 0.04 |
| $NO_2^-$ | Overall | N-mmol/h | - | - | 0.03 ± 0.05 | 0.01 ± 0.02 |
| $NH_4^+$ | Overall | N-mmol/h | 0.49 ± 0.02 | 0.47 ± 0.01 | -0.22 ± 0.02 | -0.27 ± 0.04 |
| $O_2$ | Aerobic | mmol/h | 210 | 210 | -1.0 ± 0.1 | -1.6 ± 0.2 |
| | "Overall" | mmol/h | 141 | 149 | -0.70 ± 0.07 | -1.2 ± 0.2 |
| Biomass[a] | Overall | C-mmol/h | - | - | 1.08 ± 0.09 | 1.4 ± 0.2 |
| Acetate[b] | Overall | C-mmol/h | 0.85 ± 0.02 | 1.02 ± 0.01 | -0.85 ± 0.02 | -1.02 ± 0.01 |
| Propionate[b] | Overall | C-mmol/h | 0.91 ± 0.02 | 1.09 ± 0.01 | -0.91 ± 0.02 | -1.09 ± 0.01 |
| Butyrate[b] | Overall | C-mmol/h | 0.68 ± 0.02 | 0.82 ± 0.01 | -0.69 ± 0.02 | -0.83 ± 0.01 |

[a] Calculated from the $NH_4^+$ consumption rates.
[b] Always 0 in the effluent.

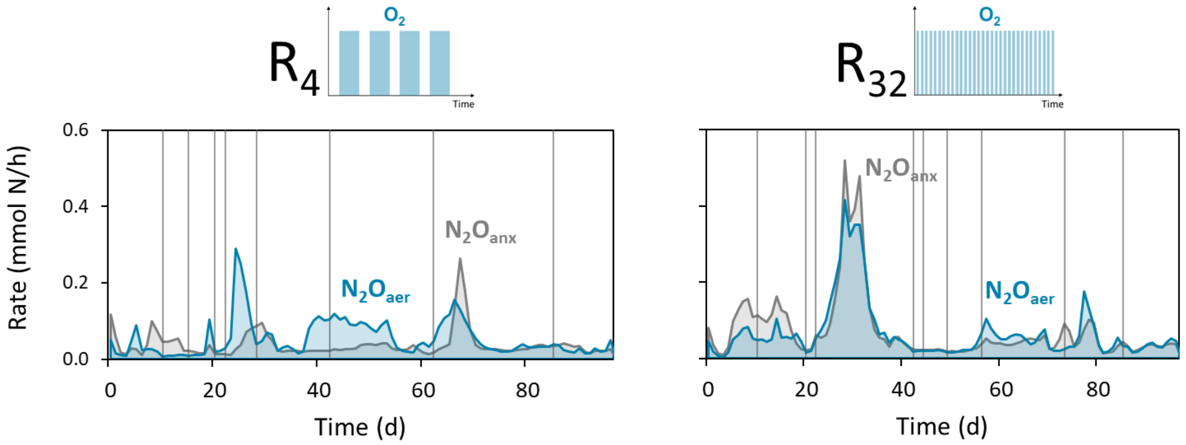


**Figure S1.** Wall-growth cleaning events (vertical lines) did not affect the profile of the aerobic (blue) and anoxic (grey) $N_2O$ emissions in the low- ($R_4$) and high-frequency ($R_{32}$) reactors.

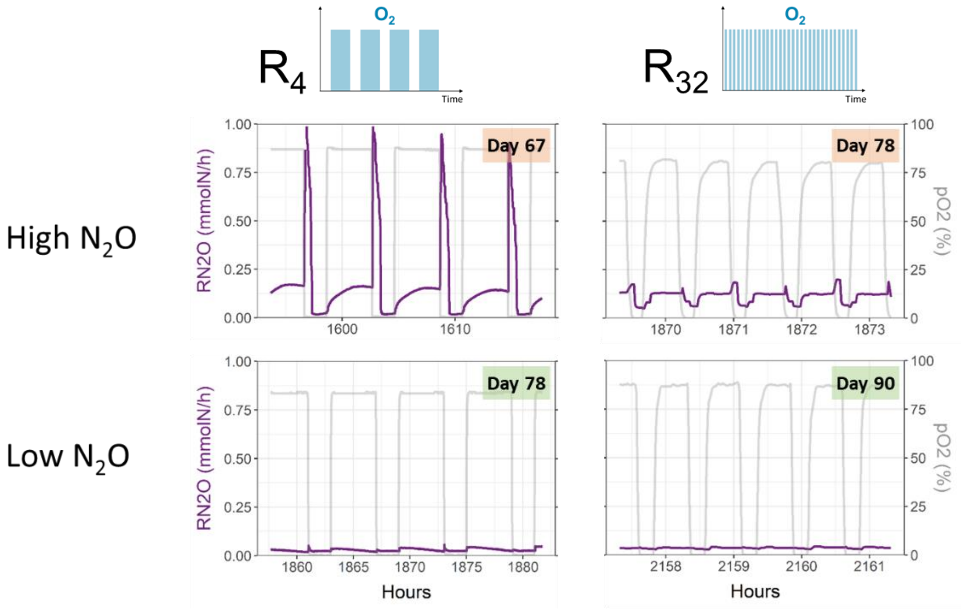


**Figure S2. $N_2O$ accumulation rates (purple, left axis) during the oxic/anoxic cycles, identifiable by the dissolved oxygen levels (grey, right axis), in the low- ($R_4$) and high-frequency ($R_{32}$) reactors.** The cycle profiles are characteristic of high (top) and low (bottom) emission periods. $N_2O$ concentrations consistently increased and plateaued during aeration. During anoxia, after the initial peak, $N_2O$ was progressively and fully consumed.

**$k_La$ determination.** The oxygen volumetric mass transfer coefficient ($k_La$) was determined to calculate the oxygen transfer rate in the aerobic phase. The $k_La$ of $R_4$ and $R_{32}$ were determined under identical conditions as the enrichments (500 rpm stirring, 400 mL/min gas flow), but with water instead of biomass. The $O_2$ transfer rates were determined by following the dissolved oxygen concentration during the sparging of air (400 mL/min) in anoxic water. The $k_La$ was obtained by fitting the integrated mass transfer equation to the dissolved $O_2$ concentration profile over time, with $C_{O2}^*$ the solubility of $O_2$ at 20 °C:

$$C_{O_2} = C^*_{O_2} \cdot (1 - e^{-k_La \cdot t}) \qquad \text{(eq. S1)}$$

The obtained $k_La$ values were 36.2 ($R_4$) and 37.0 $h^{-1}$ ($R_{32}$).

### 2. Calculation of consumption and production rates

Consumption and production rates of all dissolved and gaseous compounds were measured or estimated in the aerobic and anoxic phases, and overall (combined aerobic and anoxic). Consumption rates are negative and production rates are positive.

**Overall consumption and production rates in the liquid.** Consumption and production rates of $NO_3^-$, $NO_2^-$ ($C_{in} = 0$), $NH_4^+$ and the organic compounds acetate, propionate and butyrate ($C_{out} = 0$) were calculated from a mass balance:

$$R_i(\mathrm{mmol \cdot h^{-1}}) = F_{i,out} \cdot C_{i,out} - F_{i,in} \cdot C_{i,in} \quad \text{(eq. S2)}$$

with $R_i$ the molar rate (mmol·h$^{-1}$), $F_i$ the influent and effluent flow rates (L·h$^{-1}$), and $C_i$ the concentration of compound i (mmol·L$^{-1}$). $C_{out}$ was the average of three effluent measurements (taken at the beginning and end of the oxic phase, and end of the anoxic phase). The sample for $C_{in}$ was taken directly at the entry point of the reactors, yet differences with stock feed solution remained negligible throughout the experimental period. The flow rates were the average of the measured flow rates during the entire operation. Linear error propagation was applied to determine the standard deviation in the rates (eq. S3), using the standard deviations of $F_{in}$, $F_{out}$ and $C_{out}$ (deviation between the three measurements).

Linear error propagation of a function f dependent on multiple variables (x, y, …):

$$f(x, y, \dots): \sigma_f = \sqrt{\left(\frac{\partial f}{\partial x}\right)^2 \cdot \sigma_x^2 + \left(\frac{\partial f}{\partial y}\right)^2 \cdot \sigma_y^2 + \cdots} \quad \text{(eq. S3)}$$

with $\sigma_f$, $\sigma_x$ and $\sigma_y$ the standard deviations of f, x, and y, respectively, and $\partial f/\partial x$ and $\partial f/\partial y$ the partial derivatives of f with respect to x and y, respectively.

**Overall, aerobic and anoxic consumption and production rates in the gas phase.** A script was written in RStudio to calculate the overall, and separate oxic and anoxic $N_2O$ and $CO_2$ rates from continuous measurements recorded every minute. The molar gas flow leaving the reactor was calculated for each time point based on the constant influent volumetric gas flow rate (400 mL·min$^{-1}$) and the measured temperature and atmospheric pressure:

$$N_{gas}(\mathrm{mmol \cdot h^{-1}}) = \frac{P_{atm} \cdot F_{V,gas}}{R \cdot T} \quad \text{(eq. S4)}$$

with $N_{gas}$ the molar gas flow rate (mmol·h$^{-1}$), $P_{atm}$ the atmospheric pressure (mbar), $F_{V,gas}$ the volumetric gas flow rate, R the ideal gas constant (L·mbar·K$^{-1}$·mmol$^{-1}$) and T the reactor temperature (K). The molar gas fractions were normalized to the zero measurement before further calculations, by subtracting the corresponding value measured for the zero concentration. For each of the gases, the molar flow rates in the off-gas were calculated at each time point from measured gas fractions and the total molar gas flow rate:

$$N_{CO_2}(\mathrm{mmol \cdot h^{-1}}) = y_{CO_2} \cdot N_{gas} \quad \text{(eq. S5)}$$

The $N_2O$ rate was normalized per mole of nitrogen:

$$N_{N_2O}(\mathrm{Nmmol \cdot h^{-1}}) = 2 \cdot y_{N_2O} \cdot N_{gas} \qquad \text{(eq. S6)}$$

with $N_i$ the molar gas flow rates (mmol·h$^{-1}$), $N_{gas}$ the molar gas flow rate (mmol·h$^{-1}$) and $y_i$ the molar fractions of each compound in the off-gas. The accumulation rates at every time point were calculated with the following mass balance:

$$R_{CO_2,N_2O}\ ((\mathrm{N})\mathrm{mmol \cdot h^{-1}}) = N_{i,out} - N_{i,in} \qquad \text{(eq. S7)}$$

The average fraction of $CO_2$ in the influent air was 450 ppm. The dataset containing the rates at every minute was split in aerobic and anoxic periods, with the aerobic period defined for time points with DO > 1% (0.08 mg $O_2$·L$^{-1}$). Daily average aerobic, anoxic and overall rates were calculated with the corresponding dataset. The standard deviation of these averages was taken as the uncertainty in the rates.

**Overall and aerobic consumption and production rates of oxygen.** The $O_2$ consumption rates during the aerobic phase were calculated from the dissolved oxygen measurements during maximum aeration periods (> 20% $O_2$ in the off-gas and dissolved oxygen > 70%):

$$R_{O2}(\mathrm{mmol \cdot h^{-1}}) = k_La \cdot H_{O_2} \cdot P_{atm} \cdot y_{O_2} \cdot (1 - \mathrm{DO}) \cdot V \qquad \text{(eq. S8)}$$

with $k_La$ the experimentally measured transfer coefficient (h$^{-1}$), $H_{O2}$ the Henry coefficient for $O_2$ (0.001283 mmol·L$^{-1}$·mbar$^{-1}$), $P_{atm}$ the atmospheric pressure (mbar), $y_{O2}$ the $O_2$ molar fraction in the off-gas, DO the measured dissolved oxygen and V the broth volume (L). Daily averages were calculated and taken for further calculations. The standard deviation of these averages were taken as the uncertainty of the rates. The "overall" consumption rate of $O_2$ for further electron balancing purposes over an entire cycle was taken as the weighted average of the aerobic and anoxic (=0) rates:

$$R_i^{overall} = \frac{t_{aerobic}}{24} \cdot R_i^{aerobic} + \frac{t_{anoxic}}{24} \cdot R_i^{anoxic} \qquad \text{(eq. S9)}$$

with $t_{aerobic}$ and $t_{anoxic}$ (h) the total time in one day in which the dissolved oxygen was above or below 1%, respectively.

**Overall respiratory electron flow to nitrogen oxides and $O_2$.** The absolute and relative overall flows of electrons from organic electron donors to the electron acceptors $NO_3^-$ and $O_2$ were calculated from the overall rates considering four and five electrons for the conversion of $NO_3^-$ to $N_2O$ and $N_2$, respectively, and four electrons for the reduction of $O_2$ to $H_2O$ (

**Table S2**). $NO_3^-$ and $O_2$ were the sole electron acceptors and $NH_4^+$ fully sustained biomass growth (detailed in the following section), minimizing $NO_3^-$ assimilation. Thus, both substrates account for the entirety of the catabolic electron flow.

**Table S2.** Absolute (mmol $e^-$/h) and relative (%) overall electron flows from organic carbon to the electron acceptors $NO_3^-$ and $O_2$ in the low- ($R_4$) and high-frequency ($R_{32}$) reactors. The electron flows were calculated from the $NO_3^-$ and $O_2$ consumption and the $N_2O$ accumulation rates.

| Electron flow | # $e^-$ | $R_4$ (mmol $e^-$/h) | $R_{32}$ (mmol $e^-$/h) |
|---|---|---|---|
| $NO_3^- \rightarrow N_2O$ | 4 | 0.2 ± 0.1 | 0.2 ± 0.1 |
| $NO_3^- \rightarrow N_2$ | 5 | 3.4 ± 0.4 | 2.8 ± 0.2 |
| $O_2 \rightarrow H_2O$ | 4 | 2.8 ± 0.3 | 4.7 ± 0.6 |
| % $NO_3^-$ / Total | | 56 ± 4 % | 39 ± 4 % |
| % $O_2$ / Total | | 44 ± 4 % | 61 ± 4 % |

**Biomass production rates.** The biomass concentration was estimated from $NH_4^+$ measurements and carbon balances. In the studied system, ammonia oxidation was fully inhibited via continuous ATU addition, thus the assimilation into biomass (0.2 N-mol/C-mol) is the sole $NH_4^+$ consumption process. The biomass production rate can be calculated as follows:

$$R_X = \left| R_{NH_4^+}/0.2 \right| \qquad \text{(eq. S10)}$$

The carbon balance includes only the organic carbon substrates (acetate, propionate and butyrate), $CO_2$ and biomass, as no other products were detected in the HPLC. Therefore, the biomass production rate can also be directly calculated according to the following equation:

$$R_X\ (\mathrm{Cmmol \cdot h^{-1}}) = |R_{Ace} + R_{Pro} + R_{But} + R_{CO2}| \qquad \text{(eq. S11)}$$

For all calculations, an empirical biomass formula of $CH_{1.8}N_{0.2}O_{0.6}$ was used. The biomass concentration ($C_X$ in C-mmol·$L^{-1}$) can then be estimated from the production rate ($R_X$) and the flow rate ($F_{out}$ in L·$h^{-1}$):

$$C_X = R_X/F_{out} \qquad \text{(eq. S12)}$$

Both methods resulted in similar estimations, showing that the biomass concentration and its production rate can be determined through either one of the methods (Figure S3). The biomass rate based on $NH_4^+$ measurements was used for further calculations. Error propagation was applied to determine the standard deviation in the rates, using the standard deviations of $F_{out}$, and $NH_4^+$, organic carbon and $CO_2$ rates.

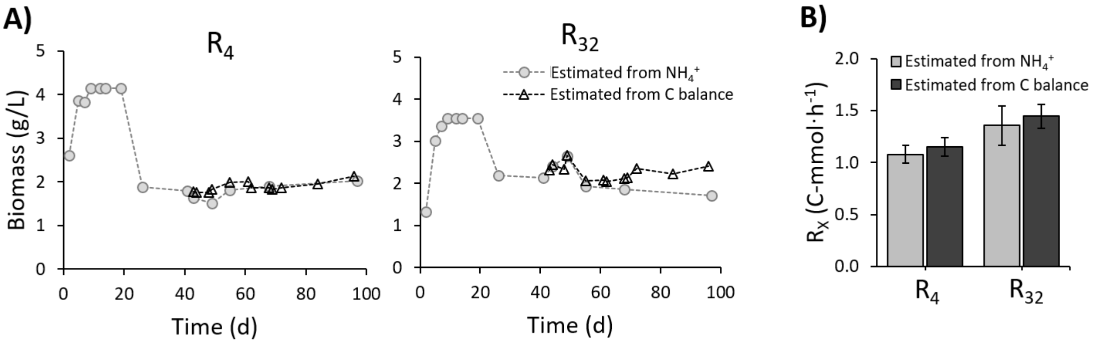


**Figure S3. Highly comparable biomass concentrations and production rates in the low- ($R_4$) and high-frequency ($R_{32}$) reactors estimated with two different methods. Panel A:** Biomass concentration over time in both the low- and high-frequency reactors, expressed as g/L. **Panel B:** average biomass production rates during the steady-state. The estimated concentrations and rates were determined from the $NH_4^+$ measurements (light grey) and the carbon balance (dark grey), i.e. the organic substrate and $CO_2$ measurements.

**Overall carbon, nitrogen and electron balances.** Mass balances were performed using the consumption and production rates averaged over the steady-state period to ensure that all substrates and products were recovered. The overall carbon balance was calculated from the consumption and production rates of acetate, propionate, butyrate, biomass (estimated from the $NH_4^+$ rates) and $CO_2$:

$$\text{C balance (\%)} = \frac{R_{in}}{R_{out}} = \frac{|R_{Ace}+R_{Pro}+R_{But}|}{R_X+R_{CO_2}} \qquad \text{(eq. S13)}$$

The nitrogen compounds involved in the nitrogen balance would be $NH_4^+$, biomass, $NO_3^-$, $NO_2^-$, NO, $N_2O$ and $N_2$. All compounds were measured (or estimated, in the case of biomass) except NO and $N_2$. NO and $NO_2^-$ accumulation was absent or negligible throughout the entire experiment, so we can assume that the missing nitrogen was recovered as $N_2$, representing full denitrification from $NO_3^-$:

$$R_{N_2} = \left|R_{NO_3^-}\right| - R_{N_2O} \qquad \text{(eq. S14)}$$

Based on all calculated and estimated rates, an electron balance was calculated.

$$e^- \text{ balance (\%)} = \frac{R_{eD}}{R_{eA}} = \frac{|4\cdot R_{Ace}+4.7\cdot R_{Pro}+5\cdot R_{But}|}{-8\cdot R_{NO_3^-}-4\cdot R_{N_2O}-3\cdot R_{N_2}-4\cdot R_{O_2}+4.2\cdot R_X} \qquad \text{(eq. S15)}$$

Uncertainty of the balances were calculated through linear error propagation from the standard deviations of the respective rates  (eq. S3).

**Table S3.** Overall carbon and electron balances over the entire steady-state period of the low- ($R_4$) and high-frequency ($R_{32}$) oxic/anoxic cycling denitrifying reactors.

| Reactor | Low-frequency | High-frequency |
|---|---|---|
| Carbon balance | 103 ± 5 % | 101 ± 9 % |
| Electrons balance | 103 ± 8 % | 100 ± 8 % |

**Estimation of the consumption and production rates in the aerobic and anoxic phases.** Separate aerobic and anoxic rates were calculated for all compounds continuously measured in the gas ($N_2O$ and $CO_2$) or liquid phase ($O_2$). In turn, grab samples for the quantification of all other compounds were less sensitive to the small concentration changes occurring during each phase, so consumption and production rates could not be determined directly. Instead, aerobic and anoxic rates were calculated from the overall mass balance and the phase-specific $N_2O$, $CO_2$, and $O_2$ rates as detailed below. In short, as the overall balances close (Table S4), all biological processes taking place in the controlled environments of the reactor are known. Also, the overall rates are the sum of the aerobic and anoxic ones weighted by their corresponding time fractions. To this end, three scenarios are considered.

- **Scenario 1: <u>no</u> aerobic conversion $NO_3^-$ to $N_2$, only to $N_2O$.** From literature, it is known that aerobic denitrification is not commonly observed in a denitrifying microbial community, at least not at a significant rate. However, in this study, significant $N_2O$ production was observed during the aerated periods. So, at first, the aerobic $NO_3^-$ consumption rate was assumed equal to the observed $N_2O$ production, excluding any $N_2$ production:

$$R_{NO_3^-}^{aerobic} = -R_{N_2O}^{aerobic} \quad \text{(eq. S16)}$$

The anoxic $NO_3^-$ consumption rate was calculated from the measured overall rate and the supposed aerobic rate, knowing that the overall consumption rate comes from a balance between the aerobic and the anoxic rates (eq. S9). The $N_2$ production rate in the anoxic phase was calculated from the $NO_3^-$ and the $N_2O$ rates (eq. S14). Similarly to $NO_3^-$, the $NH_4^+$ consumption rates could not be determined in each phase individually. Therefore, differently from the overall mass balance approach, the biomass production rates in each phase were derived from the carbon mass balances (eq. S11). The validity of this estimation was proven above (Figure S5). The $NH_4^+$ consumption rate was then estimated from the biomass production rate (eq. S10). The electron balance (eq. S15) and electron gap were then calculated for both the oxic and anoxic phases (Table S4):

$$e^- \text{ gap} = R_{eD} - R_{eA} \quad \text{(eq. S17)}$$

**Table S4.** Electron balances and gaps in the aerobic and anoxic phases of the low-frequency and high-frequency reactors, assuming the exclusive conversion of $NO_3^-$ to $N_2O$ under aerobic conditions.

| **Reactor** | **Low-frequency** | | **High-frequency** | |
|---|---|---|---|---|
| Phase | Aerobic | Anoxic | Aerobic | Anoxic |
| Electrons balance | 124 ± 9 % | 69 ± 10 % | 105 ± 7 % | 81 ± 7 % |
| Electron gap (e-mmol·$h^{-1}$) | 2.1 ± 0.7 | -5.0 ± 2.3 | 0.7 ± 0.8 | -3.2 ± 1.4 |

The electron balances do not close in either of the phases in both reactors. The balances show an underestimation of electrons accepted in the aerobic phase, and an overestimation in the anoxic phase. This suggests that more $NO_3^-$ was reduced in the aerobic phase than accounted for, while an excess $NO_3^-$ reduction was accounted for in the anoxic phase.

**Scenario 2: <u>yes</u> aerobic conversion of $NO_3^-$ to both $N_2$ and $N_2O$ - closing the electron mass balance.** From the electron balances in the previous scenario and the symmetric electron gaps of the two phases, it was hypothesized that the surplus of reduced $NO_3^-$ accounted for in the anoxic phase was actually reduced in the aerobic phase. New aerobic and anoxic $NO_3^-$ consumption rates were estimated by closing the respective electron gaps, assuming full conversion of $NO_3^-$ to $N_2$ (5 $e^-$ transfer):

$$R_{NO_3^-}^{\text{Scenario 2}} = R_{NO_3^-}^{\text{Scenario 1}} - \frac{e^-_{gap}}{5} \qquad \text{(eq. S18)}$$

Linear error propagation was applied to estimate the standard deviations of the derived rates (eq. S3). The validity of the calculations were confirmed by comparing the recalculated overall $NO_3^-$ consumption rates in scenario 2 (eq. S9) to the measured rates (Figure S4). The estimated aerobic and anoxic $NO_3^-$ consumption rates were 0.48 ± 0.14 and 1.13 ± 0.54 N-mmol/h ($R_4$) and 0.17 ± 0.17 and 1.34 ± 0.31 N-mmol/h ($R_{32}$).

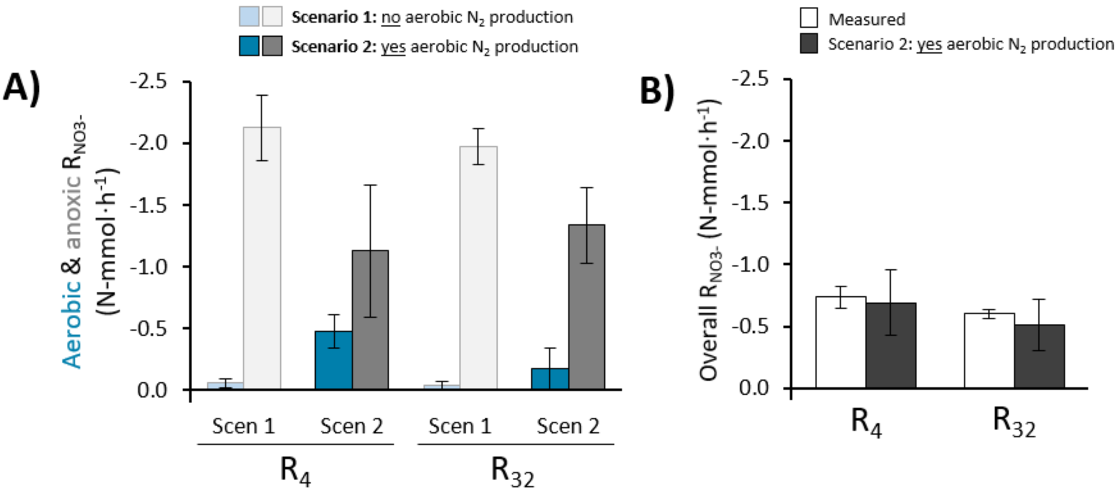


**Figure S4. Nitrate consumption rates in the aerobic (blue) and anoxic (grey) phases of the low- ($R_4$) and high-frequency ($R_{32}$) reactors. Panel A:** comparison between the predicted rates according to scenario 1 (light, assuming <u>no</u> aerobic $N_2$ production) and scenario 2 (dark, assuming <u>yes</u> aerobic $N_2$ production). **Panel B:** The measured and estimated overall nitrate consumption rates were also compared to validate the calculations.

- **Scenario 3: <u>yes</u> aerobic conversion of $NO_3^-$ to both $N_2$ and $N_2O$, and simultaneous PHA accumulation.** Cyclic conditions may select for populations accumulating storage compounds, such as polyhydroxyalkanoates (PHAs). We assessed the potential impact on the estimated aerobic $NO_3^-$ consumption rates in scenario 2 of PHA accumulation in the anoxic phase and its subsequent consumption in the aerobic period. Biomass contains 4.2 electrons per carbon, while polyhydroxybutyrate (PHB, the most common form of PHA) contains 4.5 electrons per carbon, so changes in the electron balance were minimal. Assuming that 50% of the biomass growth in the anoxic phase was actually PHA accumulation, the estimated aerobic and anoxic $NO_3^-$ consumption rates were 0.50 ± 0.14 and 1.09 ± 0.53 N-mmol/h ($R_4$) and 0.20 ± 0.18 and 1.29 ± 0.30 N-mmol/h ($R_{32}$), nearly identical to scenario 2. Therefore, our conclusions would remain unchanged even in the case of significant PHA accumulation.

### 3. Nitrification assays

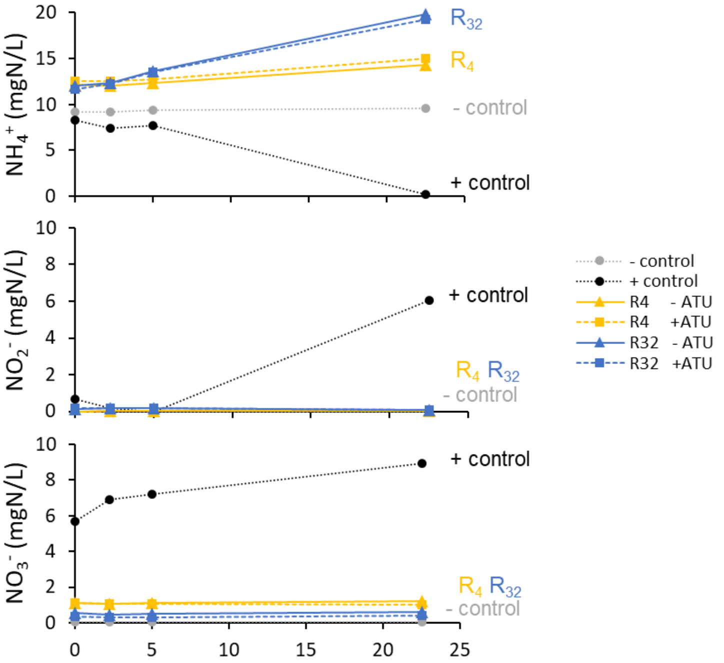


**Figure S5. Ammonium, nitrite and nitrate concentration profiles during ammonium oxidation activity tests with biomass extracted from low- ($R_4$, yellow) and high-frequency ($R_{32}$, blue) reactors, alongside a negative (water) and a positive control (nitrifying mixed culture).** Batches were performed with 10 mg $NH_4^+$-N/L, in the presence or absence of ATU.

$NH_4^+$ oxidation activity tests were performed with biomass extracted from $R_4$ and $R_{32}$, in the presence and absence of the $NH_4^+$ oxidation inhibitor ATU. A negative control replaced biomass with water and a positive control contained biomass from an enriched nitrifying microbial community. The nitrifying culture at pH 7 with a biomass concentration of 0.04 gVSS/L was enriched from activated sludge in a 2 L continuously-stirred tank reactor for 53 days, with an HRT of 4.2 days, using $NH_4^+$ as energy source (supplied at 34 $NH_4^+$-N mmol/d), bicarbonate as carbon source and $O_2$ as electron acceptor (provided as air at 500 mL/min). The nitrifying biomass was centrifuged and the pellet was resuspended in PBS buffer and added to rubber sealed bottles (filled with air) to prevent excessive evaporation. The bottles were incubated overnight in a shaker at room temperature after addition of 10 mg-N/L $NH_4^+$ to start the batches. $NH_4^+$ consumption and $NO_2^-$ and $NO_3^-$ production were observed only in the positive control. $NH_4^+$ concentrations increased in the experiments with biomass from $R_4$ and $R_{32}$, indicating biomass decay. Identical concentration profiles between experiments performed with or without ATU further confirm the absence of $NH_4^+$ oxidation activity in $R_4$ and $R_{32}$.

### 4. Metagenomics

**Table S5. Characteristics of the draft genomes recovered from $R_4$ ordered from high to low abundance (top 10 + others):** genome completeness, contamination and size, GC content, number of predicted genes, relative abundance at 68 days of operation, and taxonomic classification. Bins with a completeness lower than 70% or contamination above 10% were grouped with the unbinned portion. Low-abundant medium- and high-quality bins were grouped into "others" (grey) in the main manuscript.

| Bin | Comp(%) | Cont(%) | Genome (Mbp) | GC(%) | Genes | Abund(%) | Phylum | Class | Order | Family | Genus | Species |
|---|---|---|---|---|---|---|---|---|---|---|---|---|
| Bin1.1 | 98.96 | 0 | 5.0 | 66.1 | 4695 | 40.00 | Proteobacteria | Gammaproteobacteria | Burkholderiales | Rhodocyclaceae | *Denitromonas* | |
| Bin1.2 | 98.96 | 0.91 | 5.5 | 66.7 | 5347 | 17.37 | Proteobacteria | Alphaproteobacteria | Rhodobacterales | Rhodobacteraceae | *Wagnerdoeblera* | |
| Bin1.3 | 95.89 | 1.05 | 3.8 | 60.8 | 3400 | 7.57 | Proteobacteria | Alphaproteobacteria | Rhizobiales | Xanthobacteraceae | *Xanthobacter* | |
| Bin1.4 | 99.61 | 0 | 2.8 | 63.9 | 2582 | 3.01 | Proteobacteria | Gammaproteobacteria | Burkholderiales | Burkholderiaceae | *Brachymonas* | *Brachymonas denitrificans* |
| Bin1.5 | 99.53 | 2.24 | 3.9 | 59.4 | 3859 | 2.37 | Proteobacteria | Gammaproteobacteria | Burkholderiales | Burkholderiaceae | *Castellaniella* | |
| Bin1.6 | 98.13 | 0.29 | 2.9 | 36.3 | 2601 | 2.35 | Bacteroidota | Bacteroidia | Flavobacteriales | Flavobacteriaceae | | |
| Bin1.7 | 99.44 | 0.81 | 4.7 | 63.2 | 4678 | 1.62 | Proteobacteria | Alphaproteobacteria | Rhodobacterales | Rhodobacteraceae | *Paracoccus* | *Paracoccus sp002359815* |
| Bin1.8 | 94.12 | 0 | 3.3 | 60.5 | 2965 | 1.24 | Proteobacteria | Alphaproteobacteria | Rhizobiales | Xanthobacteraceae | *Xanthobacter* | |
| Bin1.9 | 98.48 | 3.89 | 4.5 | 62.5 | 4466 | 1.23 | Proteobacteria | Alphaproteobacteria | Rhodobacterales | Rhodobacteraceae | *Pararhodobacter* | |
| Bin1.10 | 100 | 1.67 | 4.1 | 61.7 | 3805 | 1.20 | Proteobacteria | Gammaproteobacteria | Burkholderiales | Burkholderiaceae | *Giesbergeria* | |
| Bin1.11 | 99.51 | 0.49 | 2.7 | 34.5 | 2528 | 1.01 | Bacteroidota | Bacteroidia | Chitinophagales | Chitinophagaceae | *Ginsengibacter* | |
| Bin1.12 | 97.59 | 1.09 | 3.8 | 48.1 | 3649 | 0.57 | Proteobacteria | Gammaproteobacteria | Pseudomonadales | Pseudomonadaceae | *Pseudomonas_C* | |
| Bin1.13 | 94.83 | 6.9 | 3.7 | 63.3 | 3666 | 0.46 | Proteobacteria | Gammaproteobacteria | Burkholderiales | Rhodocyclaceae | *Thauera* | |
| Bin1.14 | 97.05 | 1.14 | 4.0 | 38.9 | 3659 | 0.40 | Bacteroidota | Bacteroidia | Flavobacteriales | Flavobacteriaceae | *Aequorivita* | |
| Bin1.15 | 90.82 | 3.82 | 3.7 | 64.7 | 3856 | 0.26 | Proteobacteria | Gammaproteobacteria | Burkholderiales | Burkholderiaceae | *Giesbergeria* | |
| Bin1.16 | 81.01 | 0.55 | 2.8 | 67.2 | 3490 | 0.25 | Proteobacteria | Alphaproteobacteria | Caulobacterales | Caulobacteraceae | *Brevundimonas* | |
| Bin1.17 | 78.5 | 1.27 | 2.9 | 60 | 3425 | 0.24 | Actinobacteriota | Actinomycetia | Actinomycetales | Microbacteriaceae | *Leucobacter* | |
| Bin1.18 | 75.66 | 6.85 | 2.5 | 66.5 | 2765 | 0.23 | Proteobacteria | Gammaproteobacteria | Burkholderiales | Burkholderiaceae | *Castellaniella* | |
| Bin1.19 | 91.89 | 0.26 | 4.2 | 55.2 | 3800 | 0.10 | Proteobacteria | Gammaproteobacteria | Burkholderiales | Burkholderiaceae | *Pusillimonas_B* | *Pusillimonas_B sp013416395* |
| Bin1.20 | 87.25 | 2.93 | 3.2 | 64.7 | 3091 | 0.09 | Proteobacteria | Gammaproteobacteria | Burkholderiales | Burkholderiaceae | *Giesbergeria* | |
| Bin1.21 | 99.97 | 0.16 | 3.9 | 67.5 | 3572 | 0.08 | Proteobacteria | Gammaproteobacteria | Burkholderiales | Burkholderiaceae | *Castellaniella* | |
| Bin1.22 | 81.14 | 2.96 | 2.7 | 64.5 | 2521 | 0.08 | Proteobacteria | Gammaproteobacteria | Burkholderiales | Burkholderiaceae | *Comamonas* | |
| Bin1.23 | 79.66 | 0 | 1.3 | 25.7 | 1183 | 0.07 | Patescibacteria | JAEDAM01 | BD1-5 | UBA6164 | *UBA7396* | |
| Bin1.24 | 96.16 | 1.83 | 3.6 | 34.1 | 3183 | 0.07 | Bacteroidota | Bacteroidia | NS11-12g | UKL13-3 | *UBA6183* | |
| Bin1.25 | 91.25 | 1.21 | 3.6 | 63.5 | 3680 | 0.05 | Proteobacteria | Alphaproteobacteria | Rhodobacterales | Rhodobacteraceae | *Paracoccus* | *Paracoccus sp002359815* |
| Bin1.26 | 94.75 | 2.21 | 4.7 | 68.1 | 4687 | 0.05 | Proteobacteria | Alphaproteobacteria | Rhodobacterales | Rhodobacteraceae | *Pararhodobacter* | |
| Bin1.27 | 94.12 | 1.23 | 3.5 | 63.9 | 3394 | 0.03 | Proteobacteria | Gammaproteobacteria | Burkholderiales | Burkholderiaceae | *Giesbergeria* | *Giesbergeria suum* |
| Bin1.28 | 99.17 | 1.64 | 4.7 | 37.9 | 4055 | 0.02 | Bacteroidota | Bacteroidia | Flavobacteriales | Flavobacteriaceae | *Gelidibacter* | *Gelidibacter japonicus* |
| Bin1.29 | 86.78 | 0.99 | 2.7 | 32.7 | 2849 | 0.01 | Bacteroidota | Bacteroidia | Chitinophagales | JADIYW01 | | |
| Unbinned | | | | | | 17.99 | | | | | | |

**Table S6. Characteristics of the draft genomes recovered from $R_{32}$ ordered from high to low abundance (top 10 + others):** genome completeness, contamination and size, GC content, number of predicted genes, relative abundance at 68 days of operation, and taxonomic classification. Bins with a completeness lower than 70% or contamination above 10% were grouped with the unbinned portion. Low-abundant medium- and high-quality bins were grouped into "others" (grey) in the main manuscript.

| Bin | Comp(%) | Cont(%) | Genome (Mbp) | GC(%) | Genes | Abund(%) | Phylum | Class | Order | Family | Genus | Species |
|---|---|---|---|---|---|---|---|---|---|---|---|---|
| Bin2.1 | 98.85 | 0 | 3.2 | 59.3 | 3036 | 23.52 | Proteobacteria | Gammaproteobacteria | Burkholderiales | Burkholderiaceae | Castellaniella | |
| Bin2.2 | 99.37 | 2.4 | 3.1 | 62.2 | 2902 | 10.61 | Proteobacteria | Gammaproteobacteria | Burkholderiales | Burkholderiaceae | Castellaniella | |
| Bin2.3 | 98.81 | 1.76 | 4.3 | 63 | 3964 | 7.32 | Proteobacteria | Gammaproteobacteria | Burkholderiales | Rhodocyclaceae | Thauera | |
| Bin2.4 | 98.29 | 0.4 | 3.6 | 36.9 | 3206 | 5.02 | Bacteroidota | Bacteroidia | Flavobacteriales | Flavobacteriaceae | Aequorivita | |
| Bin2.5 | 99.41 | 0.47 | 3.9 | 68.9 | 3479 | 2.67 | Proteobacteria | Gammaproteobacteria | Burkholderiales | Burkholderiaceae | Castellaniella | Castellaniella defragrans |
| Bin2.6 | 93.95 | 2.25 | 3.1 | 61.8 | 2958 | 2.13 | Proteobacteria | Gammaproteobacteria | Burkholderiales | Burkholderiaceae | Comamonas | |
| Bin2.7 | 98.99 | 0.91 | 3.8 | 59.2 | 3671 | 1.65 | Proteobacteria | Alphaproteobacteria | Rhodobacterales | Rhodobacteraceae | Pseudorhodobacter | |
| Bin2.8 | 97.73 | 3.66 | 5.0 | 63 | 5008 | 1.52 | Proteobacteria | Alphaproteobacteria | Rhodobacterales | Rhodobacteraceae | Paracoccus | Paracoccus sp002359815 |
| Bin2.9 | 96.7 | 2.46 | 3.0 | 37.2 | 2922 | 1.44 | Bacteroidota | Bacteroidia | Flavobacteriales | Flavobacteriaceae | | |
| Bin2.10 | 96.36 | 4.9 | 3.9 | 68.8 | 3558 | 0.95 | Proteobacteria | Gammaproteobacteria | Burkholderiales | Burkholderiaceae | Melaminivora_A | |
| Bin2.11 | 80.22 | 0 | 1.3 | 26.6 | 1250 | 6.53 | Patescibacteria | JAEDAM01 | BD1-5 | UBA6164 | UBA7396 | UBA7396 sp002470645 |
| Bin2.12 | 94.69 | 7.47 | 3.8 | 62.7 | 3530 | 3.76 | Proteobacteria | Gammaproteobacteria | Burkholderiales | Burkholderiaceae | Comamonas | Comamonas sp019104825 |
| Bin2.13 | 97.7 | 7.04 | 4.1 | 48.4 | 3929 | 2.18 | Proteobacteria | Gammaproteobacteria | Pseudomonadales | Pseudomonadaceae | Pseudomonas_C | |
| Bin2.14 | 88.51 | 2.61 | 3.1 | 65.9 | 3002 | 0.95 | Proteobacteria | Gammaproteobacteria | Xanthomonadales | Xanthomonadaceae | Stenotrophomonas | |
| Bin2.15 | 77.08 | 1.63 | 1.8 | 43.1 | 1964 | 0.89 | Proteobacteria | Alphaproteobacteria | Paracaedibacterales | Paracaedibacteraceae | | |
| Bin2.16 | 92.88 | 5.36 | 3.7 | 64.8 | 3865 | 0.81 | Proteobacteria | Gammaproteobacteria | Burkholderiales | Burkholderiaceae | Giesbergeria | |
| Bin2.17 | 81.35 | 0 | 1.2 | 25.9 | 1111 | 0.35 | Patescibacteria | JAEDAM01 | BD1-5 | UBA6164 | UBA7396 | |
| Bin2.18 | 98 | 0.03 | 3.3 | 68.8 | 3065 | 0.33 | Proteobacteria | Gammaproteobacteria | Burkholderiales | Burkholderiaceae | Castellaniella | |
| Bin2.19 | 90.25 | 3.15 | 3.1 | 64.1 | 2759 | 0.27 | Proteobacteria | Gammaproteobacteria | Burkholderiales | Burkholderiaceae | Comamonas | |
| Bin2.20 | 92.18 | 2.05 | 3.5 | 64.6 | 3616 | 0.20 | Proteobacteria | Alphaproteobacteria | Rhizobiales | Devosiaceae | Devosia | |
| Bin2.21 | 97.98 | 1.84 | 3.9 | 38.7 | 3378 | 0.16 | Bacteroidota | Bacteroidia | Flavobacteriales | Flavobacteriaceae | Aequorivita | |
| Bin2.22 | 97.27 | 3.95 | 4.5 | 62.3 | 4359 | 0.10 | Proteobacteria | Gammaproteobacteria | Burkholderiales | Rhodocyclaceae | Azoarcus_C | |
| Bin2.23 | 93.02 | 1.94 | 3.8 | 60.2 | 3838 | 0.05 | Proteobacteria | Alphaproteobacteria | Rhizobiales | Rhizobiaceae | Hoeflea | |
| Bin2.24 | 78.42 | 3.71 | 3.2 | 67.1 | 3605 | 0.04 | Proteobacteria | Alphaproteobacteria | Sphingomonadales | Sphingomonadaceae | Sphingopyxis | Sphingopyxis granuli |
| Bin2.25 | 99.29 | 0.5 | 3.2 | 64.5 | 3015 | 0.04 | Proteobacteria | Gammaproteobacteria | Burkholderiales | Burkholderiaceae | Castellaniella | |
| Bin2.26 | 92.51 | 1 | 3.1 | 68.1 | 2884 | 0.04 | Proteobacteria | Gammaproteobacteria | Burkholderiales | Burkholderiaceae | Comamonas_C | Comamonas_C sp002894305 |
| Bin2.27 | 77.62 | 2.16 | 2.6 | 64 | 2734 | 0.04 | Proteobacteria | Gammaproteobacteria | Burkholderiales | Burkholderiaceae | Giesbergeria | |
| Bin2.28 | 77.69 | 0.98 | 2.0 | 61.5 | 2259 | 0.03 | Actinobacteriota | Actinomycetia | Actinomycetales | Microbacteriaceae | Leucobacter | |
| Bin2.29 | 88.65 | 2.51 | 2.9 | 48 | 3153 | 0.02 | Proteobacteria | Gammaproteobacteria | Pseudomonadales | Pseudomonadaceae | Pseudomonas_C | |
| Unbinned | | | | | | 26.38 | | | | | | |

**Table S7.** Reference KO-numbers of the genes from the nitrogen metabolism.

| KO ID | Gene | Description | Pathway |
|---|---|---|---|
| K00362 | nirB | nitrite reductase (NADH) large subunit [EC:1.7.1.15] | DNRA |
| K00363 | nirD | nitrite reductase (NADH) small subunit [EC:1.7.1.15] | |
| K03385 | nrfA | nitrite reductase (cytochrome c-552) [EC:1.7.2.2] | |
| K15876 | nrfH | cytochrome c nitrite reductase small subunit | |
| K00370 | narG, narZ, nxrA | nitrate reductase / nitrite oxidoreductase, alpha subunit [EC:1.7.5.1 1.7.99.-] | Denitrification |
| K00371 | narH, narY, nxrB | nitrate reductase / nitrite oxidoreductase, beta subunit [EC:1.7.5.1 1.7.99.-] | |
| K00373 | narJ, narW | chaperone | |
| K00374 | narI, narV | nitrate reductase gamma subunit [EC:1.7.5.1 1.7.99.-] | |
| K07673 | narX | nitrate/nitrite sensor | |
| K07684 | narL | response regulator | |
| K02575 | NRT, narK, nrtP, nasA | MFS transporter, NNP family, nitrate/nitrite transporter | |
| K02567 | napA | nitrate reductase (cytochrome) [EC:1.9.6.1] | |
| K02568 | napB | nitrate reductase (cytochrome), electron transfer subunit | |
| K02570 | napD | nitrate reductase (cytochrome) | |
| K02571 | napE | nitrate reductase (cytochrome) | |
| K00367 | narB | ferredoxin-nitrate reductase | |
| K10850 | narT | putative nitrate transporter | |
| K15576 | nrtA, nasF, cynA | nitrate/nitrite transport system substrate-binding protein | |
| K15577 | nrtB, nasE, cynB | nitrate/nitrite transport system permease protein | |
| K15578 | nrtC, nasD | nitrate/nitrite transport system ATP-binding protein [EC:7.3.2.4] | |
| K15579 | nrtD, cynD | nitrate/nitrite transport system ATP-binding protein | |
| K21563 | dnr | CRP/FNR family transcriptional regulator, dissimilatory nitrate respiration regulator | |
| K01420 | fnr | CRP/FNR family transcriptional regulator, anaerobic regulatory protein | |
| K00368 | nirK | nitrite reductase (NO-forming) [EC:1.7.2.1] | |
| K15864 | nirS | nitrite reductase (NO-forming) / hydroxylamine reductase [EC:1.7.2.1 1.7.99.1] | |
| K04561 | norB | nitric oxide reductase subunit B [EC:1.7.2.5] | |
| K02305 | norC | nitric oxide reductase subunit C | |
| KnorZ | norZ | quinol-dependent nitric oxide reductase | |
| K02448 | norD | nitric oxide reductase D protein | |
| K02164 | norE | nitric oxide reductase E protein | |
| K04747 | norF | nitric oxide reductase F protein | |
| K04748 | norQ | nitric oxide reductase Q protein | |
| K12266 | treg | transcription regulator | |
| K13771 | trep | NO-sensitive transcription repressor | |
| K00376 | nosZ I | nitrous-oxide reductase [EC:1.7.2.4] | |
| KnosZII | nosZ II | nitrous-oxide reductase [EC:1.7.2.4] | |
| K19339 | nosR | nitrous-oxide reductase transcriptional regulator | |
| K19342 | nosL | copper chaperone | |
| K07218 | nosD | accessory protein | |
| K10944 | amoA | methane/ammonia monooxygenase subunit A [EC:1.14.18.3 1.14.99.39] | Nitrification |
| K10945 | amoB | methane/ammonia monooxygenase subunit B | |
| K10946 | amoC | methane/ammonia monooxygenase subunit C | |
| K10535 | hao | hydroxylamine dehydrogenase [EC:1.7.2.6] | |

**Table S8**. Reference KO-numbers of the genes from the respiratory chain.

| KO ID | Gene | Description | Reaction type |
|---|---|---|---|
| K00330 | nuoA | NADH-quinone oxidoreductase subunit A [EC:7.1.1.2] | |
| K00331 | nuoB | NADH-quinone oxidoreductase subunit B [EC:7.1.1.2] | |
| K00332 | nuoC | NADH-quinone oxidoreductase subunit C [EC:7.1.1.2] | |
| K00333 | nuoD | NADH-quinone oxidoreductase subunit D [EC:7.1.1.2] | |
| K00334 | nuoE | NADH-quinone oxidoreductase subunit E [EC:7.1.1.2] | |
| K00335 | nuoF | NADH-quinone oxidoreductase subunit F [EC:7.1.1.2] | |
| K00336 | nuoG | NADH-quinone oxidoreductase subunit G [EC:7.1.1.2] | |
| K00337 | nuoH | NADH-quinone oxidoreductase subunit H [EC:7.1.1.2] | NADH oxidation |
| K00338 | nuoI | NADH-quinone oxidoreductase subunit I [EC:7.1.1.2] | |
| K00339 | nuoJ | NADH-quinone oxidoreductase subunit J [EC:7.1.1.2] | |
| K00340 | nuoK | NADH-quinone oxidoreductase subunit K [EC:7.1.1.2] | |
| K00341 | nuoL | NADH-quinone oxidoreductase subunit L [EC:7.1.1.2] | |
| K00342 | nuoM | NADH-quinone oxidoreductase subunit M [EC:7.1.1.2] | |
| K00343 | nuoN | NADH-quinone oxidoreductase subunit N [EC:7.1.1.2] | |
| K03885 | ndh | NADH:quinone reductase (non-electrogenic) [EC:1.6.5.9] | |
| K00411 | petA | ubiquinol-cytochrome c reductase iron-sulfur subunit [EC:7.1.1.8] | |
| K00412 | petB | ubiquinol-cytochrome c reductase cytochrome b subunit | |
| K00413 | petC | ubiquinol-cytochrome c reductase cytochrome c1 subunit | |
| K00410 | fbcH | ubiquinol-cytochrome c reductase cytochrome b/c1 subunit | Cyt c reduction |
| K03890 | qcrA | ubiquinol-cytochrome c reductase iron-sulfur subunit | |
| K03891 | qcrB | ubiquinol-cytochrome c reductase cytochrome b subunit | |
| K03889 | qcrC | ubiquinol-cytochrome c reductase cytochrome c subunit | |
| K02274 | coxA,ctaD | cytochrome $aa_3$ oxidase subunit I [EC:7.1.1.9] | |
| K02275 | coxB,ctaC | cytochrome $aa_3$ oxidase subunit II [EC:7.1.1.9] | |
| K02276 | coxC,ctaE | cytochrome $aa_3$ oxidase subunit III [EC:7.1.1.9] | |
| K02277 | coxD,ctaF | cytochrome $aa_3$ oxidase subunit IV [EC:7.1.1.9] | |
| K00404 | ccoN | cytochrome $cbb_3$ oxidase subunit I [EC:7.1.1.9] | |
| K00405 | ccoO | cytochrome $cbb_3$ oxidase subunit II | |
| K15862 | ccoNO | cytochrome $cbb_3$ oxidase subunit I/II [EC:7.1.1.9] | |
| K00407 | ccoQ | cytochrome $cbb_3$ oxidase subunit IV | |
| K00406 | ccoP | cytochrome $cbb_3$ oxidase subunit III | $O_2$ reduction |
| K02297 | cyoA | cytochrome *bo*3 ubiquinol oxidase subunit I [EC:7.1.1.3] | |
| K02298 | cyoB | cytochrome *bo*3 ubiquinol oxidase subunit II [EC:7.1.1.3] | |
| K02299 | cyoC | cytochrome *bo*3 ubiquinol oxidase subunit III | |
| K02300 | cyoD | cytochrome *bo*3 ubiquinol oxidase subunit IV | |
| K00425 | cydA | cytochrome *bd* ubiquinol oxidase subunit I [EC:7.1.1.7] | |
| K00426 | cydB | cytochrome *bd* ubiquinol oxidase subunit II [EC:7.1.1.7] | |
| K00424 | cydX | cytochrome *bd* ubiquinol oxidase subunit X [EC:7.1.1.7] | |
| K08738 | CYC | cytochrome c | Cytochrome c |
| K02111 | atpA | F-type H+/Na+-transporting ATPase subunit alpha [EC:7.1.2.2 7.2.2.1] | |
| K02108 | atpB | F-type H+-transporting ATPase subunit a | |
| K02114 | atpC | F-type H+-transporting ATPase subunit epsilon | |
| K02112 | atpD | F-type H+/Na+-transporting ATPase subunit beta [EC:7.1.2.2 7.2.2.1] | ATP synthesis |
| K02110 | atpE | F-type H+-transporting ATPase subunit c | |
| K02109 | atpF | F-type H+-transporting ATPase subunit b | |
| K02115 | atpG | F-type H+-transporting ATPase subunit gamma | |
| K02113 | atpH | F-type H+-transporting ATPase subunit delta | |

**Table S9.** Reference KO-numbers of the genes from the ROS-protection pathway.

| KO ID | Gene | Description | Reaction type |
|---|---|---|---|
| K04565 | SOD1 | superoxide dismutase, Cu-Zn family [EC:1.15.1.1] | $O_2^-$ → $H_2O_2$ |
| K04564 | SOD2 | superoxide dismutase, Fe-Mn family [EC:1.15.1.1] | |
| K03781 | katE, catB, srpA | catalase [EC:1.11.1.6] | |
| K07217 | Mn-cat | Mn-catalase | |
| K03782 | katG | catalase-peroxidase [EC:1.11.1.21] | |
| K00428 | ccp | cytochrome c peroxidase [EC:1.11.1.5] | |
| K00430 | px | peroxidase | |
| K11065 | tpx | thiol peroxidase | $H_2O_2$ → $H_2O$ |
| K00432 | gpx, btuE, bsaA | glutathione peroxidase | |
| K05910 | npr | NADH peroxidase | |
| K14171 | ahp1 | alkyl hydroperoxide reductase | |
| K03386 | ahpC | alkyl hydroperoxide reductase | |

### 5. Heatmaps with gene presence and protein expression

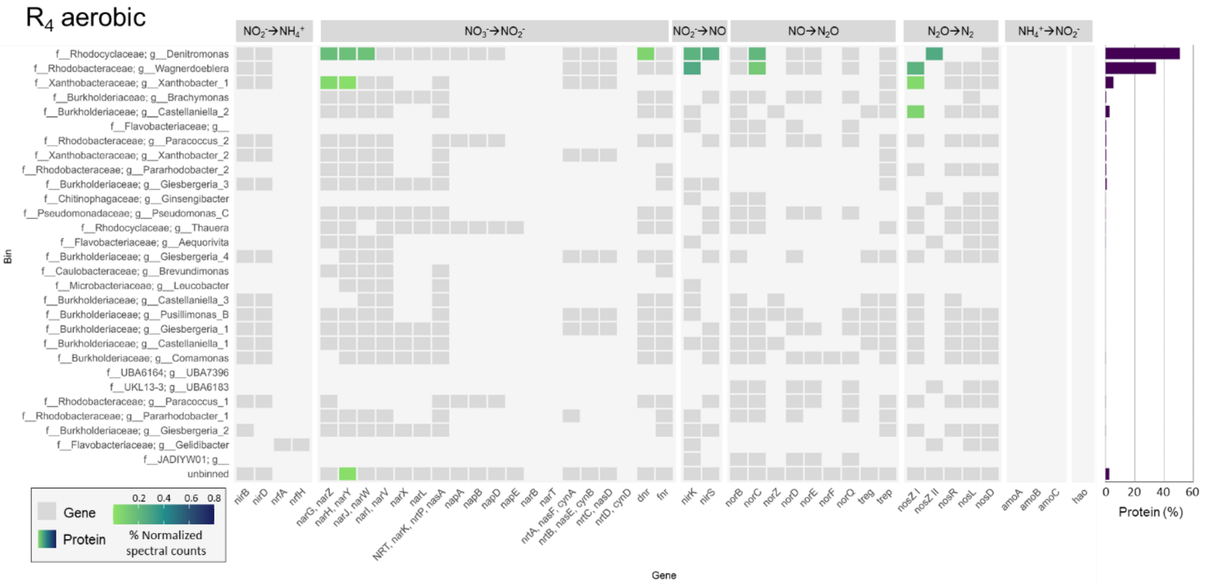


**Figure S6.** Heatmap with gene presence (grey) and protein expression (coloured) of the nitrogen metabolism, represented as relative abundance of the total proteome, of all MAGs (ordered from high to low abundance in the metagenome) at the end of the aerobic phase of R4. **Right bar charts:** total relative abundance of each MAG in the metaproteome.

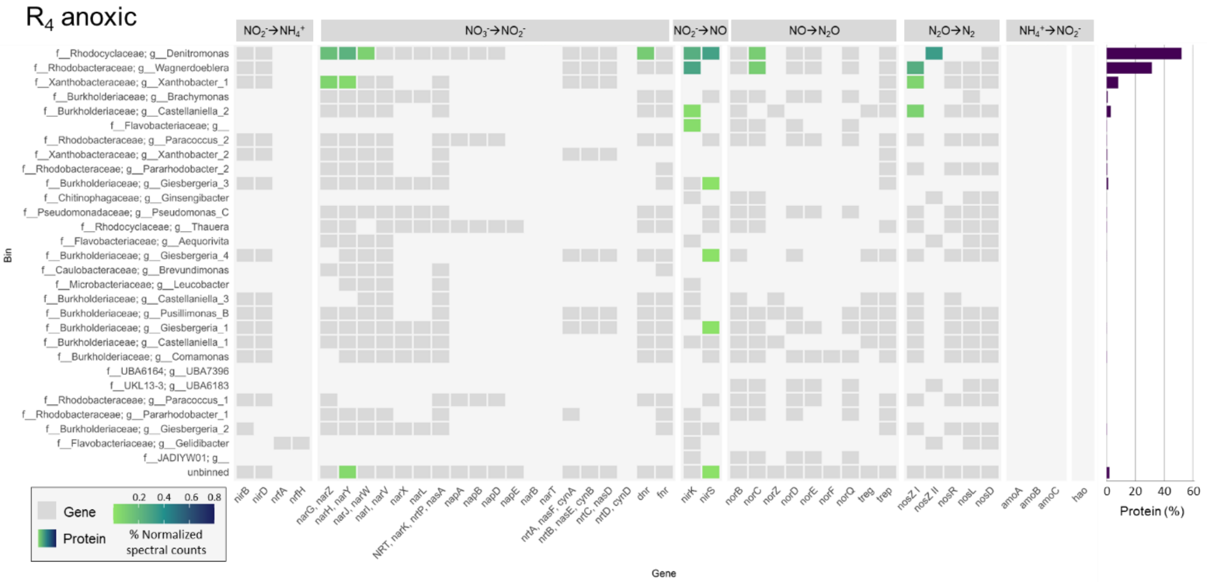


**Figure S7.** Heatmap with gene presence (grey) and protein expression (coloured) of the nitrogen metabolism, represented as relative abundance of the total proteome, of all MAGs (ordered from high to low abundance in the metagenome) at the end of the anoxic phase of R4. **Right bar charts:** total relative abundance of each MAG in the metaproteome.

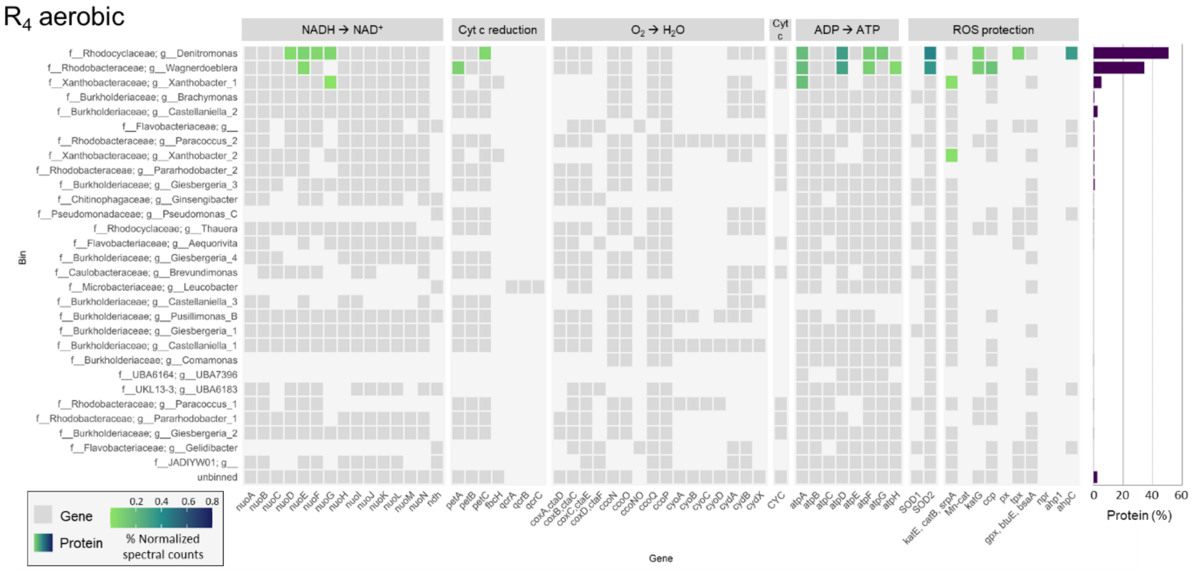


**Figure S8.** Heatmap with gene presence (grey) and protein expression (coloured) of the respiratory chain and ROS-protection pathway, represented as relative abundance of the total proteome, of all MAGs (ordered from high to low abundance in the metagenome) at the end of the aerobic phase of $R_4$. **Right bar charts:** total relative abundance of each MAG in the metaproteome.

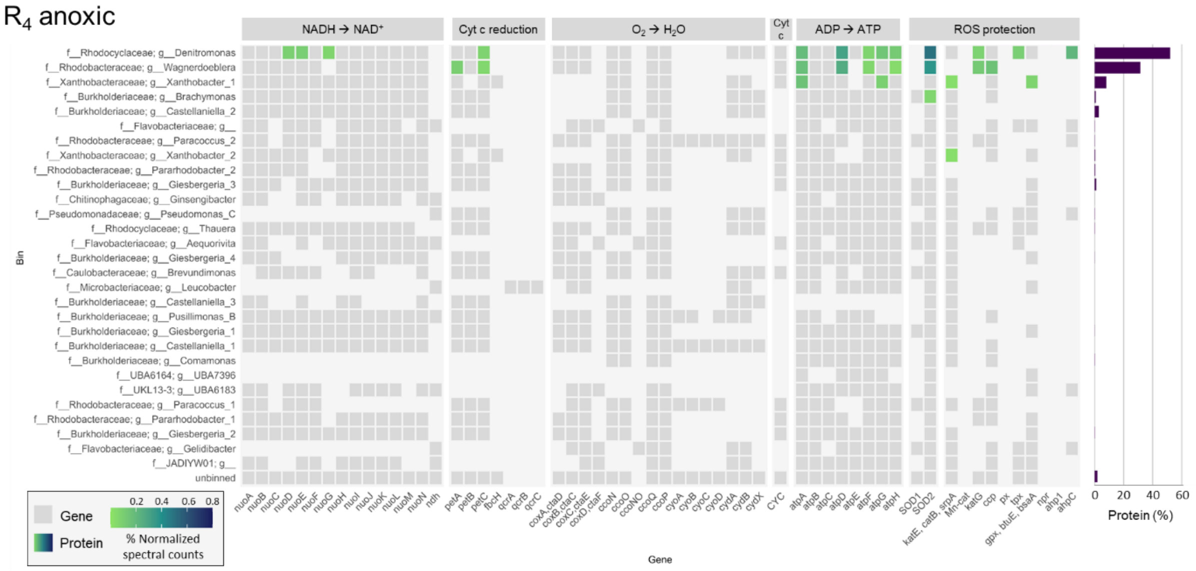


**Figure S9.** Heatmap with gene presence (grey) and protein expression (coloured) of respiratory chain and ROS-protection pathway, represented as relative abundance of the total proteome, of all MAGs (ordered from high to low abundance in the metagenome) at the end of the anoxic phase of $R_4$. **Right bar charts:** total relative abundance of each MAG in the metaproteome.

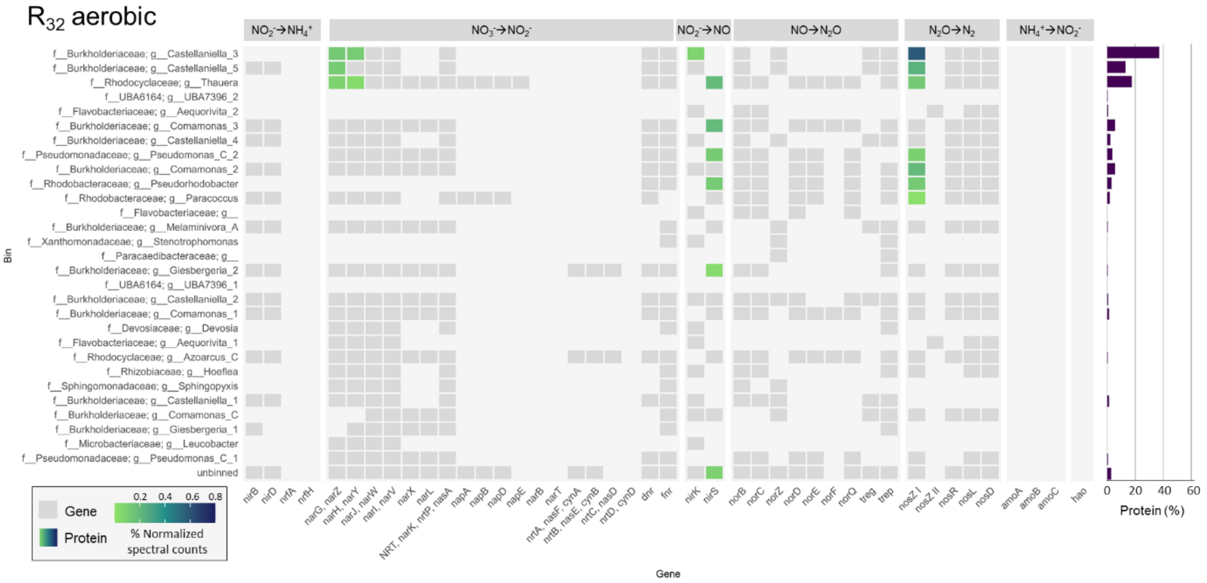


**Figure S10.** Heatmap with gene presence (grey) and protein expression (coloured) of the nitrogen metabolism, represented as relative abundance of the total proteome, of all MAGs (ordered from high to low abundance in the metagenome) at the end of the aerobic phase of $R_{32}$. **Right bar charts:** total relative abundance of each MAG in the metaproteome.

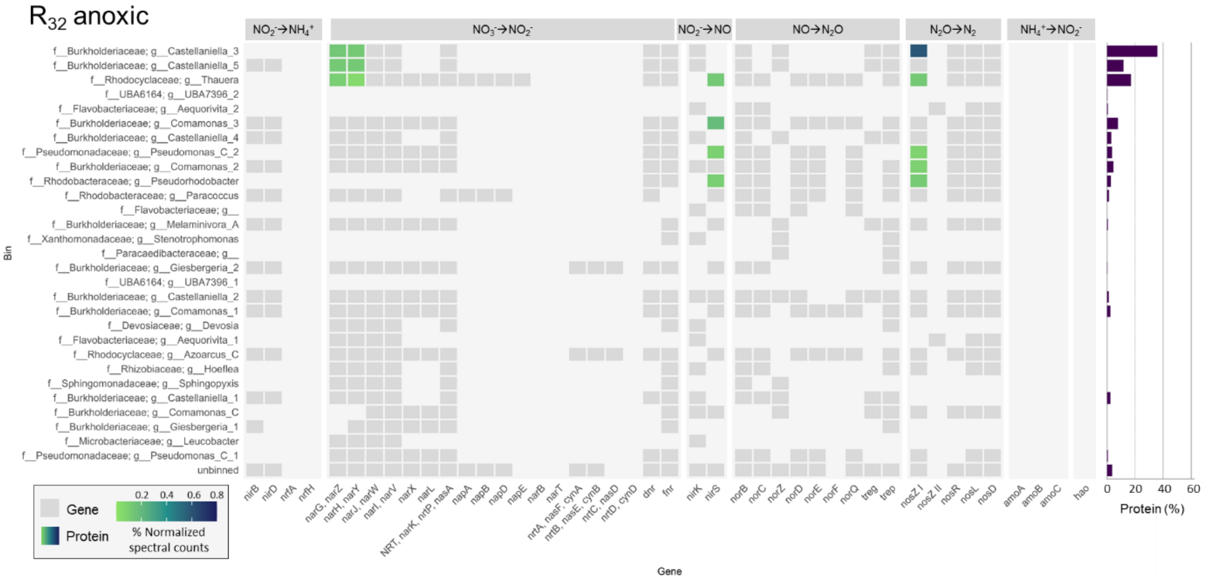


**Figure S11.** Heatmap with gene presence (grey) and protein expression (coloured) of the nitrogen metabolism, represented as relative abundance of the total proteome, of all MAGs (ordered from high to low abundance in the metagenome) at the end of the anoxic phase of $R_{32}$. **Right bar charts:** total relative abundance of each MAG in the metaproteome.

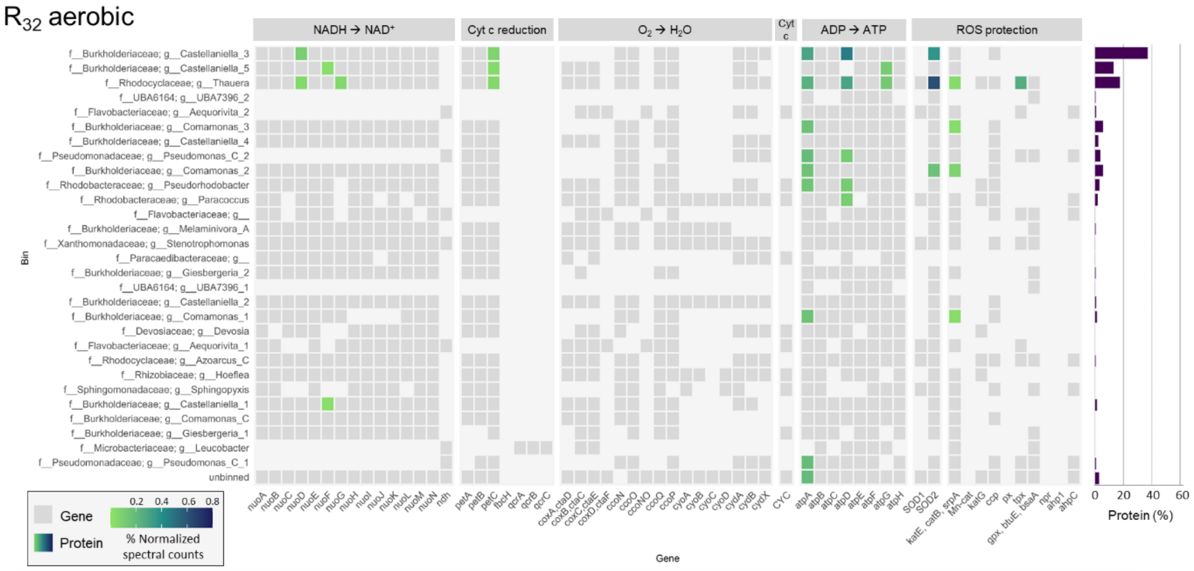


**Figure S12.** Heatmap with gene presence (grey) and protein expression (coloured) of respiratory chain and ROS-protection pathway, represented as relative abundance of the total proteome, of all MAGs (ordered from high to low abundance in the metagenome) at the end of the aerobic phase of $R_{32}$. **Right bar charts:** total relative abundance of each MAG in the metaproteome.

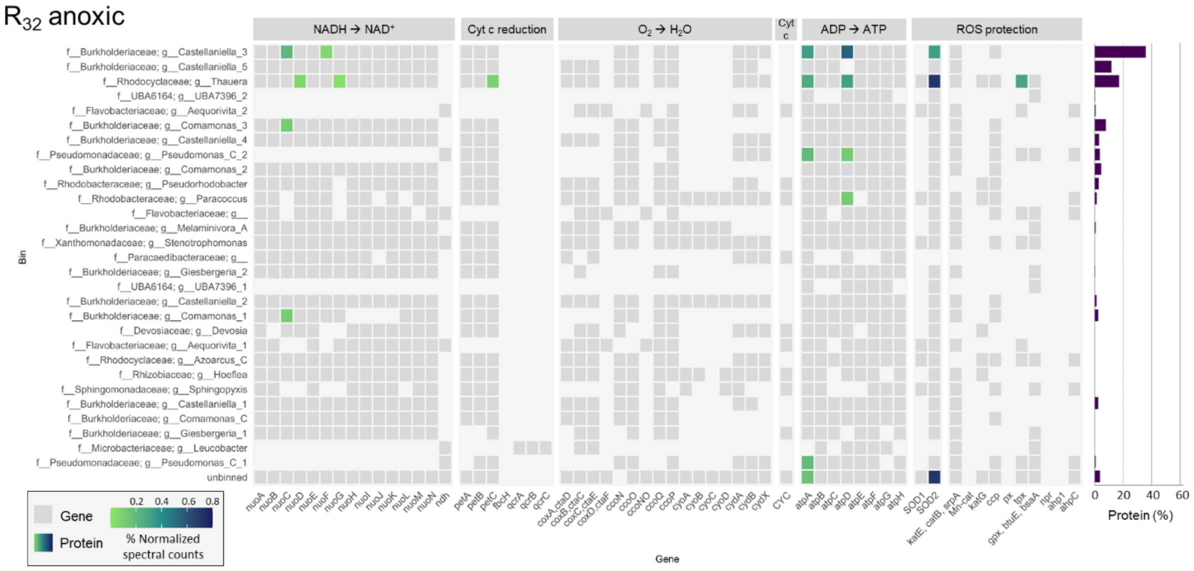


**Figure S13.** Heatmap with gene presence (grey) and protein expression (coloured) of respiratory chain and ROS-protection pathway, represented as relative abundance of the total proteome, of all MAGs (ordered from high to low abundance in the metagenome) at the end of the anoxic phase of $R_{32}$. **Right bar charts:** total relative abundance of each MAG in the metaproteome.

### 6. Oxic-anoxic cycling in 5 reference Dutch WWTPs

The exposure frequency of activated sludge to oxic-anoxic cycles in wastewater treatment plants (WWTPs) cannot exactly be determined, but estimations were made for different WWTPs using flow rates and tank volumes. The hydraulic residence time in each of the tanks was determined (Figure S14): anaerobic (no $O_2$, no $NO_x$), anoxic (no $O_2$), facultative (can function as anoxic or aerobic tank, according to the treatment needs), and aerobic. The sludge residence time in the anoxic tanks varied between 11 and 142 minutes, while this was 13-155 min for the aerobic zones. The biomass that passes through the settler experiences approximately one oxic-anoxic transition per day, which is equivalent to 15-42 transitions per sludge retention time (SRT, equivalent to the cell generation time, normally 15-20 days in a WWTP). If cells remain in a recycling loop between the aerobic and anoxic zones they can experience up to 9-36 transitions per day, i.e. 132-756 switches per SRT (Table S10). Similarly to the activated sludge in the WWTPs, the biomass in our reactors experienced 4 ($R_4$) and 32 ($R_{32}$) oxic-anoxic transitions per day, equalling 8 and 64 transitions within one SRT. Experiments with even higher frequency of oxic-anoxic transitions within one SRT should be performed to assess the extent of aerobic denitrification in the highest frequency ranges observed in WWTPs.

**Table S10.** Number of oxic-anoxic transitions experienced by the biomass in our reactors and in five different WWTP configurations (Figure S14), in one day and within one sludge retention time (SRT).

| System | SRT (d) | Cycles/day | Cycles/SRT |
|---|---|---|---|
| $R_4$ | 2 | 4 | 8 |
| $R_{32}$ | 2 | 32 | 64 |
| WWTP A | 15 | 1 - 35 | 15 - 315 |
| WWTP B | 21 | 2 - 36 | 42 - 756 |
| WWTP C | 22 | 1 - 19 | 22 - 418 |
| WWTP D | 15 | 1 - 14 | 15 - 210 |
| WWTP E | 15 | 1 - 9 | 15 - 135 |

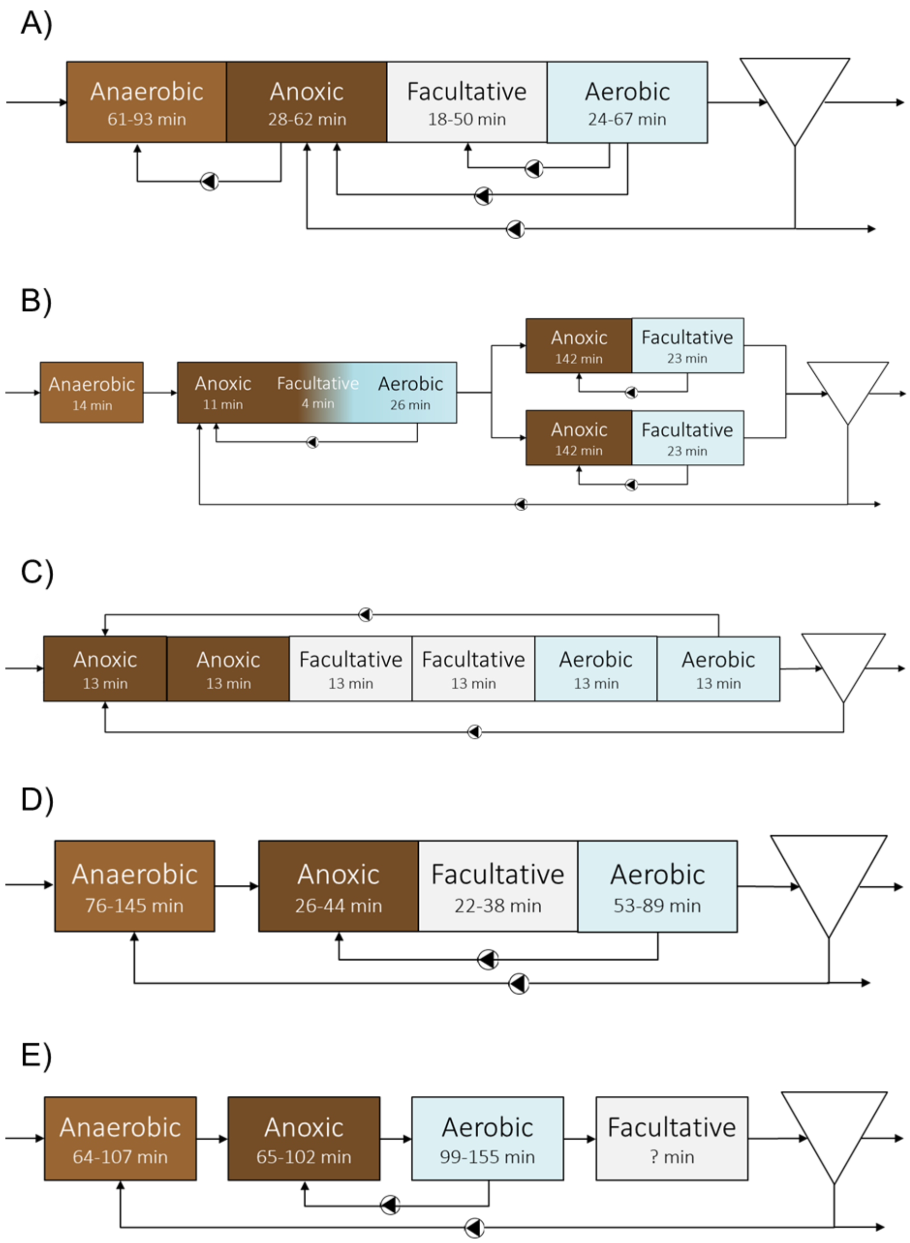


**Figure S14. Hydraulic residence time in tanks with different conditions in five different Dutch WWTPs, representing the time that the sludge experiences those conditions.** The anaerobic tanks do not contain $O_2$ nor nitrogen oxides, the anoxic tanks have no oxygen, the aerobic tanks are aerated with air and the facultative tanks can function either as anoxic or aerobic tanks.
